# Tracking propagating cortical activity in MEG/EEG with a bilinear state-space model

**DOI:** 10.64898/2026.09.02.748579

**Authors:** Anna Kubiak, Nikita Fedosov, Alex Ossadtchi

## Abstract

Magnetoencephalography (MEG) and electroencephalography (EEG) are ideal for studying macroscopic neural dynamics, but non-invasive tracking of cortical traveling waves remains a major methodological challenge. Traditional inverse solutions assume spatiotemporal separability, restricting sources to fixed spatial topographies. Consequently, they struggle to capture the continuous spatial migration of cortical traveling waves and often misinterpret phase-locked static sources as spurious propagation. To address this fundamental limitation, we propose a dynamic state-space framework that explicitly accommodates the spatiotemporal inseparability of propagating neural activity. Our approach models the sensor signal as a bilinear combination of two states that evolve together: a fast, narrowband stochastic oscillator carrying the electrical time course, and a slowly evolving spatial topography that drifts through a data-driven singular value decomposition subspace. Both the rhythmic electrical time series and the migrating source trajectory track jointly via an Unscented Kalman Filter. We evaluated the method on realistically simulated MEG data and empirical resting-state MEG and EEG recordings targeting the occipital alpha rhythm. In simulations, the approach accurately recovered electrical time courses and spatial trajectories across varying signal-to-noise ratios, spatial envelope velocities up to 0.1 m/s, and distinct cortical geometries (calcarine and central sulci), significantly outperforming traditional minimum norm estimation and dipole fitting. Crucially, the model resists fabricating spurious propagation trajectories when presented with stationary, coherent dipoles. Application to empirical MEG and EEG recordings of the occipital alpha rhythm yields anatomically plausible, temporally cohesive propagation paths that explain significantly more sensor-level variance than static baselines. By embedding the evolving source geometry directly into the inverse solution, this framework provides a robust, proof-of-concept tool for the non-invasive investigation of macroscopic propagating brain dynamics.

## 1 Introduction

Magnetoencephalography (MEG) and electroencephalography (EEG) are non-invasive neuroimaging methods ideally suited for studying the fine features of spatio-temporal dynamics exhibited by neural circuits. However, extracting functionally meaningful information from these recordings critically depends on the approach used to solve the ill-posed inverse problem. Traditional analysis methods in cognitive neuroscience often assume the spatiotemporal separability of cortical processes — an assumption deeply rooted in Donders’ paradigm formulated in the late 19th century [1]. According to this paradigm, neural activity can be represented as a linear combination of static sources, allowing cognitive processes to be studied through subtraction or averaging methods. However, modern neuroscience [2] and advancements in multichannel recording technologies [3, 4] have revealed new aspects of neural activity that severely violate the assumptions of this classical theory. This is particularly evident in the case of cortical traveling waves (TWs) that continuously propagate across the cortical surface [5].

Cortical TWs manifest a multiscale, complex neural dynamic emerging from the activity of millions of interconnected cortical neurons. As they spread, individual TWs can overlap and interact, creating complex spatiotemporal patterns that vary in speed [6, 7], scale [8, 9], shape [10, 11], origin [9, 7, 12], propagation direction [13, 10], and functional significance [13, 11, 9, 14]. By producing local and global brain states, TWs are thought to play a key role in information processing: they have been observed across the sensory and motor cortices of various species [15, 10], in the human neocortex across multiple frequency bands [5, 14, 16, 6], and as slow waves during sleep [17]. Yet, the bulk of this evidence still derives from invasive measurements. Reliably observing TWs non-invasively represents a major methodological challenge, which imposes a serious limitation on the broader study of these phenomena. To help address this problem, in this paper, we develop and evaluate a proof-of-concept approach for non-invasive tracking of a single dominant propagating source.

It is essential to recognize that “traveling waves” in brain activity represent the general capacity of neuronal tissue to sustain spatially propagating patterns, rather than classical waves governed by standard wave equations. The implication for source modeling is that the magnetic fields and electric potentials recorded by distant sensors depend not only on the temporal dynamics of the source but also on its dynamically and concurrently changing geometric properties, such as the location and orientation of the equivalent current dipole (ECD). As neural activation propagates, the ECD geometry evolves along anatomically constrained trajectories, modulating the sensor-level timeseries in complex ways. Consequently, a prior built directly on the classical wave equation [18] is unlikely to accurately constrain the spatial dynamics of the underlying generators. This motivates a shift from sensor-level descriptions toward inverse models, where the spatial pattern of the source is itself allowed to evolve in time.

Standard source reconstruction approaches — such as dipole fitting [19], including the rotating dipole model [20], and distributed source modeling like minimum norm estimation [21] and beamforming [22] — are fundamentally designed to localize stationary or quasi-stationary sources. As such, these methods cannot accurately capture the trajectory of spatially non-stationary propagating activity. Modern instrumentation alone cannot gracefully resolve this limitation. While non-cryogenic optically pumped magnetometers [23, 24] and ultra-high-density EEG systems [25] provide an unprecedented amount of information about the electromagnetic field changes in proximity to the cortex, the inverse problem remains severely ill-posed for fundamental reasons. Overcoming this for moving sources requires regularization strategies that explicitly encode priors on the expected *spatial* as well as temporal dynamics of the underlying generators. To capture cortical TWs, a separable, time-invariant topography is no longer an adequate model.

This fact is further sharpened by a second, more insidious failure mode. As recently demonstrated [26], a pair of coherent static dipoles oscillating with a specific phase lag can generate sensor-level spatiotemporal patterns that are practically indistinguishable from cortical TWs. Traditional linear localization methods fail to resolve this ambiguity, acting as poorly tuned estimators that misinterpret phase differences as spatial propagation. Non-invasive TW research is therefore doubly fragile: separable inverse methods cannot *track* genuine propagation, and the same methods can *manufacture* the appearance of propagation from static coupled sources. Both failures share a single root cause — the absence of an explicit model of how the source topography itself evolves in time.

Several approaches do move beyond static reconstruction by introducing temporal dynamics into the inverse problem. State-space and Kalman-filtering formulations recursively constrain the source estimate through a temporal prior [27, 28], and oscillation-decomposition models explicitly capture narrow-band rhythmic dynamics [29]. These methods, however, typically retain a fixed spatial model. For instance, the dMAP-EM algorithm [27] enforces local smoothness through a static spatial coupling matrix, while the topography attributed to each source remains time-invariant. Consequently, the spatial migration that *defines* the spatial focus of the traveling activation is never part of the estimated state, leaving a critical methodological gap exactly where non-separable cortical dynamics reside.

To address this, we present a state-space approach that explicitly accommodates the temporally evolving geometry of an active neural population. At the core of the method is the treatment of the source topography as a time-varying latent state, expanded in a data-driven SVD subspace and tracked jointly with the electrical time series using an Unscented Kalman Filter. This joint dependence is consistent with the principle of spatiotemporal inseparability [2]. Throughout, we operate at the macroscopic spatial scale accessible to MEG/EEG rather than at the neuronal-population scale probed by invasive recordings. The method rests on a single structural assumption: that the source topography migrates on a slower timescale than the carrier oscillation. This assumption delimits the regime in which the approach is expected to apply. In the present study, we map out this regime empirically, characterizing the method across spatial envelope propagation velocities spanning 0.01–0.5 m/s and identifying the point at which the timescale separation ceases to hold; the occipital alpha rhythm serves as our validation case [16, 14].

Specifically, the main contributions and advantages of our proposed framework are:

- **Nonlinear Dynamic Observation Model:** We formulate a bilinear measurement model that separates the fast oscillatory electrical activity (modeled as a frequency-modulated process) from the slower spatial migration of the underlying source(modeled using an autoregressive model).
- **Data-Driven Spatial Subspace:** While other dynamic approaches, such as the dMAP-EM algorithm [27], enforce local smoothness via a fixed spatial coupling matrix, our state-space model treats the source topography itself as part of the dynamic state. We express the instantaneous spatial topography as a time-varying linear combination of abstract eigen-topographies extracted from the data matrix via SVD. Treating these combination coefficients as autoregressive processes themselves imposes the expected temporal scale of the spatial changes, furnishes natural spatial smoothness, and offers the flexibility to track true wave propagation.
- **Behavior under a known confound:** In a controlled simulation of two stationary, phase-locked dipoles — a configuration known to produce wave-like patterns at the sensor level — the model did not produce a propagating trajectory and instead returned a compact, stationary localization cluster. We show this as a qualitative property of our solution that needs to be quantified in the future to enable discrimination between TW and static activation patterns using the non-invasive data.
- **Nonlinear State Estimation:** To efficiently invert the proposed bilinear measurement model, we formulate a novel application of the Unscented Kalman Filtering (UKF) framework [30, 31, 32]. This provides a practical means of jointly estimating the spatial trajectory and the electrical time series without requiring analytic Jacobians. We note that the standard optimality guaranties of the linear Kalman filter do not carry over to this nonlinear setting.

We assess the behavior of the proposed approach using realistically simulated and experimental EEG/MEG datasets. Concretely, we (i) benchmark the method on simulations with realistic leadfield perturbations against minimum-norm, dipole-fitting, and RAP-MUSIC [33, 34] reconstructions, and characterize its accuracy across envelope propagation velocities from 0.01 to 0.5 m/s, as well as its robustness to varying signal-to-noise ratios (SNR); (ii) examine, in a controlled two-dipole scenario, whether it separates genuine propagation from static coherent sources; and (iii) apply it to real MEG and EEG datasets of the occipital alpha rhythm [35], recovering smooth, anatomically plausible source trajectories. Therefore, we illustrate that a state-space model treating the source topography as a latent variable is estimable in practice from MEG and EEG data and that, under the tested simulated conditions, allows for the recovery of the electrical time course and accurately locating the cortical propagation trajectory.

## 2 Materials and Methods

### 2.1 Traditional Observation Equation (Measurement Model)

The fundamental relationship between the recorded magnetic or electric signals and their neuronal sources [36] is mathematically expressed as

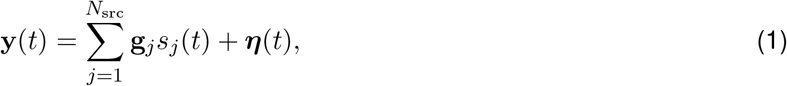

where the vector 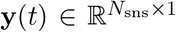 represents the set of signal values recorded by magnetic or voltage sensors at time *t*, and where *N*_sns_ corresponds to the number of sensors. Here, the summation is performed over all *j* from 1 to *N*_src_, where *N*_src_ corresponds to the number of relevant sources. The term *s*_*j*_(*t*) denotes the amplitude of the *j*-th source activity, which is unknown and varies over time. The vector 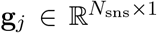 is the topography of the *j*-th source, i.e., a column of the *forward model matrix* that characterizes how a unit-amplitude neural source at a given *j*-th location contributes to the measured signals at each sensor. In other words, the elements of **g**_*j*_ quantify the sensitivity of each sensor to the activity of the j-th source. While the computation of the full leadfield matrix **G** is a well-established numerical task — solved using Maxwell’s equations under quasi-static approximations and informed by individual MRI-based anatomical models — identifying the specific columns **g**_*j*_ relevant to a given data is non-trivial. This challenge arises because the precise locations and orientations of the underlying neuronal sources are a priori unknown. Consequently, determining which spatial topographies contribute to the measured signal **y**(*t*) constitutes the core of the electromagnetic inverse problem. Finally, 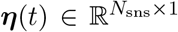 represents the noise vector, which encapsulates external environmental disturbances, sensor-specific artifacts, and ongoing brain activity unrelated to the phenomenon under study. Throughout this paper, time and space are treated as discrete. For notational convenience, explicit sample indices are omitted.

In its most general form, the full forward model incorporates a vast number of potential source locations, often reaching tens of thousands. This high dimensionality relative to the number of sensors renders the inverse problem of reconstructing neuronal activity severely ill-posed and underdetermined. To overcome this, it is necessary to incorporate prior information regarding source properties, which ensures the physical plausibility and structural consistency of the solution. Such priors allow the problem to be reformulated in a well-posed manner, facilitating a unique and stable estimation. This is typically achieved through two primary strategies: first, by introducing regularization (such as Tikhonov or *ℓ*_1_-norm penalties) to invert the full leadfield matrix, thereby balancing data fidelity with the chosen source prior in a distributed solution. Alternatively, one may assume the existence of only a limited number of focal sources, which simplifies the task to identifying a parsimonious set of source locations and their corresponding time series.

While both approaches have their merits, they generally fail to capture the non-stationary dynamics inherent in cortical TWs. Accurate modeling of such phenomena necessitates the explicit representation of spatiotemporal dependencies to account for the continuous evolution of spatial patterns — a capability largely absent in traditional “separable” frameworks, which typically assume time-invariant spatial topographies. This limitation is further compounded by the fact that the spatial progression of a wave and the local electrical dynamics represent distinct physiological processes of the underlying neuronal populations. Consequently, effective modeling requires decoupled dynamic priors: one to characterize the spatial displacement (the trajectory of the wave) and another to describe the time-varying electrical signals generated at each successive location.

### 2.2 Dynamic Observation Model

As mentioned earlier, the assumption of *spatiotemporal separability* implies that the cortical activity *F*(*r, t*) can be decomposed into a linear combination of fixed spatial patterns, each modulated by a corresponding temporal activation series. For a single component, this is expressed as the product of a time-varying amplitude *S*(*t*) and a purely spatial topography *H*(*r*). In contrast, *spatiotemporal non-separability* refers to a state or process where the spatial and temporal components cannot be decomposed into independent functions; that is:

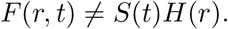

In our study, this behavior is captured by allowing the activation topography itself to evolve over time, effectively becoming a function *H*(*r, t*) of both space and time.

To integrate this non-separable behavior into the observation model defined in equation (1), we introduce an explicit temporal dependence into the topography vectors **g**_*j*_(*t*) for each source *j* = 1, …, *N*_src_:

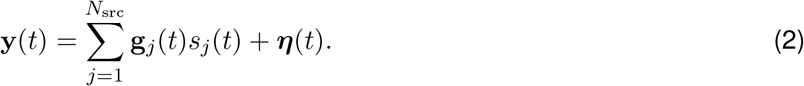

In our simulations and subsequent data analysis, we focus on scenarios dominated by a single primary source of interest (e.g., the occipital alpha rhythm), setting *N*_src_ = 1. Although our mathematical framework generalizes to multi-source scenarios, this study focuses on the fundamental case of a single source, providing a baseline for evaluating the precision of spatiotemporal tracking.

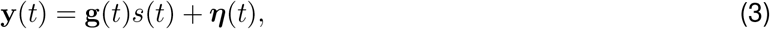

where *s*(*t*) represents the source amplitude — capturing the *local electrical dynamics* — and **g**(*t*) denotes the *time-varying topography* of the migrating source. This formulation effectively decouples the signal’s intrinsic oscillation from its spatial progression across the cortex.

This formulation, however, poses a significant challenge: without further constraints, the temporal evolution of the spatial topography **g**(*t*) would be underdetermined and physically unconstrained. To ensure that the estimated topography remains within a physically plausible and data-driven subspace, we represent the time-varying source vector as a linear combination of spatial basis vectors:

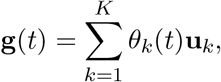

where 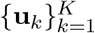 form a basis that constrains the topography to a low-dimensional subspace. The time-varying projection coefficients *θ*_*k*_(*t*) determine the instantaneous contribution of each basis vector to the topography and, consequently, dictate the spatial weighting of these components within the observed data **y**(*t*). We refer to these as *dynamic spatial coefficients*, as they capture the evolution of the source’s spatial pattern as it manifests in the recorded signal **y**(*t*).

Consequently, we propose an alternative to the classical observation model that explicitly incorporates these dynamic properties:

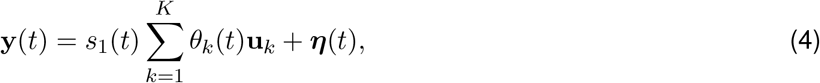

where *s*_1_(*t*) is the electrical component (source amplitude), and ***η***(*t*) represents white Gaussian measurement noise, which is assumed to follow a multivariate normal distribution ***η***(*t*) ∼ *N* (**0, R**). In this framework, *θ*_*k*_(*t*) and *s*_1_(*t*) are time-varying scalars.

The basis vectors 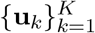 are derived from the Singular Value Decomposition (SVD) of the analyzed MEG/EEG data segment. Specifically, we utilize the matrix 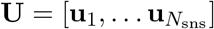, composed of left singular vectors that capture the predominant spatial patterns in the recording [33]. The measurement space spanned by these singular vectors represents linear combinations of the recorded sensor signals and encompasses both components related to the neural sources of interest and those associated with noise and measurement artifacts. This partitioning of the measurement space distinguishes between the *signal subspace* and the *noise-only subspace* — a concept formally introduced by Mosher et al. [33].

By retaining only the first *K* left singular vectors, we focus on the signal subspace, effectively performing dimensionality reduction and suppressing irrelevant activity. Crucially, while traditional static frameworks often associate each singular vector within the signal subspace with a distinct, independent neural source, our dynamical representation treats these *K* components as a basis. We argue that the spatial migration of a single neural source naturally gives rise to multiple SVD components and can be represented as a time-varying linear combination of these. Thus, instead of seeking a one-to-one mapping between sources and components, we use the first *K* singular vectors — which we refer to as *eigen-topographies* — to capture the full range of spatial variability inherent in a single moving source. In practice, *K* is selected to ensure that the set of eigen-topographies accounts for at least 95% of the total variance in the MEG/EEG data.

This approach allows us to model the spatiotemporal dynamics via a bilinear decomposition, where the observed signal is factorized into time-varying electrical and spatial components. To formalize the evolution of these components, we subsequently employ a state transition model.

### 2.3 The State Transition Model

Neural signals encompass diverse types of activity, including background noise, evoked responses, and induced activity. The latter consists of multispectral rhythmic oscillations, commonly referred to as brain rhythms. Our framework focuses on these rhythmic components, as they represent the most prevalent and ubiquitous forms of macroscopic brain activity. These rhythms are exemplified by the occipital alpha rhythm (8–12 Hz), which is typically induced by eye closure and is associated with a state of relaxed wakefulness, and the sensorimotor mu-rhythm (9–13 Hz), which reflects analogous dynamics within the cortical motor system. Other functionally significant oscillations include the delta (0.5–4 Hz), theta (4–8 Hz), beta (12–30 Hz), and gamma (30–100 Hz) rhythms. In this paper, we focus primarily on the alpha rhythm as a representative case; however, the proposed framework is generalizable and can be readily adapted to any of the aforementioned rhythmic activities by adjusting the model’s central frequency *f*_*c*_ in the transition matrix.

It has been shown [29, 37] that the characteristic peak around 10 Hz in the EEG/MEG spectrum can be modeled as a narrowband stochastic process. Specifically, the alpha rhythm is viewed not as a pure sine wave, but as an oscillation where the instantaneous phase and amplitude are subject to random perturbations. Building on this framework, we describe the source dynamics using the following state-transition equation:

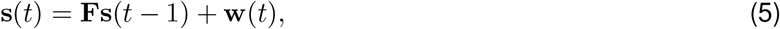

where **s**(*t*) = [*s*_1_(*t*), *s*_2_(*t*)]^*T*^ represents the analytical signal in the state space, with *s*_1_(*t*) capturing the oscillatory activity of the brain source and *s*_2_(*t*) serving as its quadrature (imaginary) component for phase estimation. The transition matrix **F** is defined as

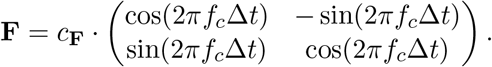

This matrix governs the deterministic part of the dynamics, representing a rotation by an angle 0 ≤ 2*πf*_*c*_Δ*t* ≤ *π* at each time step, where Δ*t* denotes a sampling interval of the analyzed data and *f*_*c*_ is the central frequency of the rhythm in Hz. The parameter 0 *< c*_**F**_ *<* 1 acts as a damping coefficient that controls the bandwidth and smoothness of the signal envelope. The process noise **w**(*t*) ∼ *N* (**0, Q**) introduces stochastic perturbations to the state, accounting for model uncertainty and influencing random external factors. In this formulation, the interaction between the rotation matrix and the process noise effectively models random fluctuations in the instantaneous phase, which account for the characteristic frequency-modulated appearance of the alpha rhythm.

The evolution of the spatial component is expected to occur on a slower timescale compared to the fluctuations of the electrical oscillatory activity. This separation of timescales is consistent with the physiological nature of TWs, where the spatial progression across the cortex is typically more gradual than the underlying carrier frequency. In the absence of a specific prior model for the wave trajectory, we make a Markov assumption regarding its dynamics. Specifically, we model the spatial coefficients as a first-order stationary autoregressive (AR(1)) process:

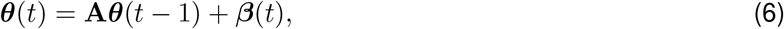

where ***θ***(*t*) = [*θ*_1_(*t*), …, *θ*_*k*_(*t*), …, *θ*_*K*_(*t*)]^*T*^ is the vector of spatial coefficients. The state transition matrix **A** is defined as **A** = *c*_**A**_ **I**, where **I** is the identity matrix, and the scalar parameter 0 *< c*_**A**_ *<* 1 ensures the stability and smoothness of the spatial trajectory within the eigen-topography subspace. The process noise ***β***(*t*) ∼ *N* (**0, B**) accounts for stochastic deviations from the predicted path, allowing the model to adapt to non-linearities or unpredictable changes in the wave’s progression.

Combining models (5) and (6) yields the following complete state-space representation:

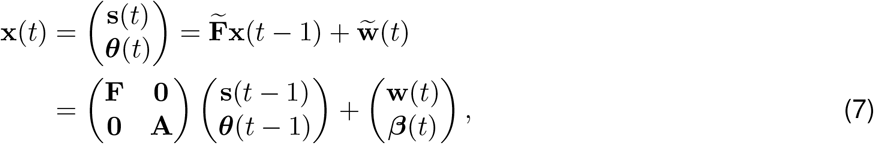

where **x**(*t*) = [*s*_1_(*t*), *s*_2_(*t*), *θ*_1_(*t*), …, *θ*_*K*_(*t*)]^*T*^ denotes the *joint state vector* comprising the unknown variables of interest. The process noise follows a multivariate normal distribution 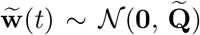, where the covariance matrix 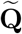 is a block diagonal matrix consisting of **Q** and **B**.

To estimate the latent variables *s*_1_(*t*), *s*_2_(*t*), and ***θ***(*t*) from the data, we combine the dynamic observation model (4) with the state-transition model (7) and employ the Kalman filter algorithm.

### 2.4 The Kalman Filter

The Kalman filter [38] is a recursive algorithm designed to estimate the state vector of a dynamical system with unknown variables by optimally combining prior knowledge of the system’s behavior with sequential sensor measurements.

The Kalman filter operates by employing two key models: the *process model*, which predicts the system’s dynamics, and the *observation model*, which provides a mapping between measured sensor data and the hidden variables of interest. This algorithm effectively integrates the process model predictions with observed data, ensuring robustness in the presence of noise and uncertainty. Importantly, the filter not only provides a point estimate of the system state but also quantifies the uncertainty of this estimate through a covariance matrix, allowing for a probabilistic interpretation of the system’s state at each time step. Moreover, the estimates obtained using the Kalman filter are optimal in “the minimum mean square error” sense.

Most versions of the Kalman filter assume discretization in the time domain, where each iteration consists of a *prediction phase* and an *update phase*. First, the algorithm provides the distribution of the *predicted state estimate*, characterized by the mean 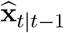 and covariance **P**_*t*|*t*−1_, based on the process model and the state estimate obtained at the previous iteration. Then, the predicted state estimate is updated using the observation model and the set of measurements obtained at the current time step. The final *updated state estimate* is *estimate*, state characterized by the mean 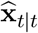 and covariance **P**_*t*|*t*_.

The working principle of the algorithm (schematically illustrated in Figure 1) can be viewed as the combination of information from two distinct sources, accounting for their respective uncertainty and noise. Mathematically, at each time step, this process corresponds to the statistical fusion of two Gaussian distributions: one associated with the prediction and the other with the measurements.

**Figure 1.**
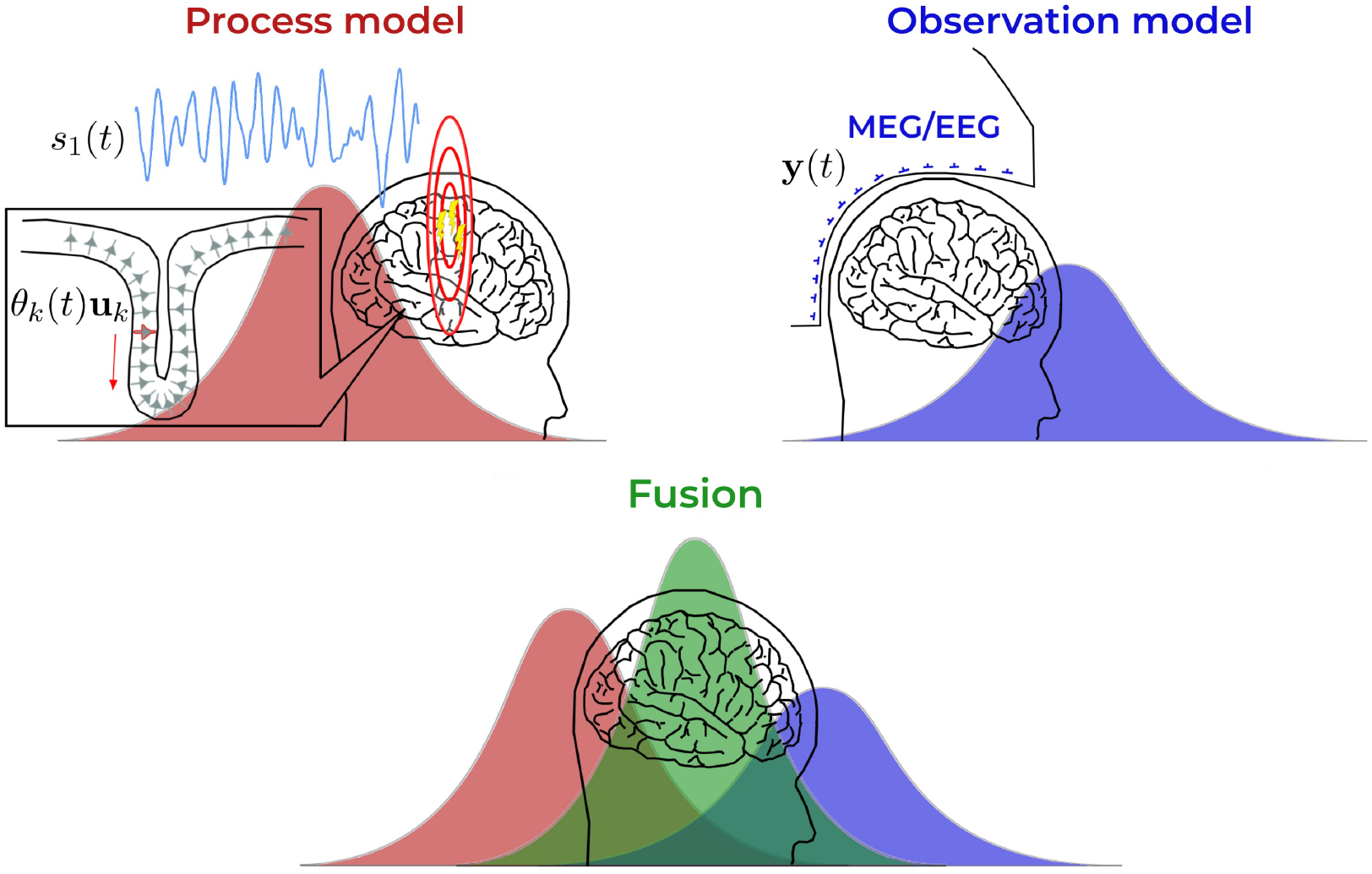
Schematic illustration of the Kalman filter implementation used in this study for estimating neural dynamics. The optimal state estimate is obtained by combining information from two sources: the *process model* and the *observation model*. The process model provides a prediction of the current system state based on the previous one. In this study, it describes the temporal dynamics of alpha oscillations (equation (5)) and the slowly varying spatial component (equation (6)). The observation model defines how the measured data relate to the state variables, with MEG/EEG signals serving as observations. The algorithm performs a statistical fusion of two Gaussian distributions: one representing the prediction (red) and the other corresponding to the observed measurements (blue). At each time step, the Kalman filter determines the relative contribution of each information source while accounting for uncertainty (noise covariance matrices). This results in an optimal estimate of the system’s state (green Gaussian).

In this study, equations (7) and (4) serve as the *process model* and the *observation model*, respectively, within the Kalman filter framework. Since the observation model (4) is *nonlinear* with respect to the state vector variables *s*_1_(*t*) and *θ*_*k*_(*t*), a nonlinear variant of the Kalman filter — the Unscented Kalman filter (UKF; [30, 31]) — is employed to handle the estimation problem.

#### 2.4.1 The Unscented Kalman Filter

The UKF is particularly effective when at least one of the system’s models is nonlinear, rendering the traditional linear Kalman filter inapplicable. To handle such nonlinearity, the UKF employs a deterministic sampling technique known as the unscented transformation (UT) to select a small set of weighted sample points, called sigma points, around the mean. These sigma points are strategically chosen to capture the mean and covariance of the system state distribution, ensuring an accurate approximation of its propagation through nonlinear transformations.

Compared to the Extended Kalman Filter (EKF), which relies on first-order linearization and requires the explicit computation of Jacobians, the UKF provides higher accuracy in many practical scenarios and does not require the system functions to be analytically differentiable [30]. Another alternative, particle filters or the sequential Monte Carlo method, can handle extreme nonlinearities and non-Gaussian noise; however, they typically require a large number of particles to achieve comparable accuracy, leading to prohibitive computational costs [39]. In contrast, the UKF offers a favorable trade-off between estimation accuracy and computational efficiency, producing reliable results with a minimal number of sigma points. Figure 2 provides a conceptual illustration of these advantages, motivating the choice of the UKF for the present work.

**Figure 2.**
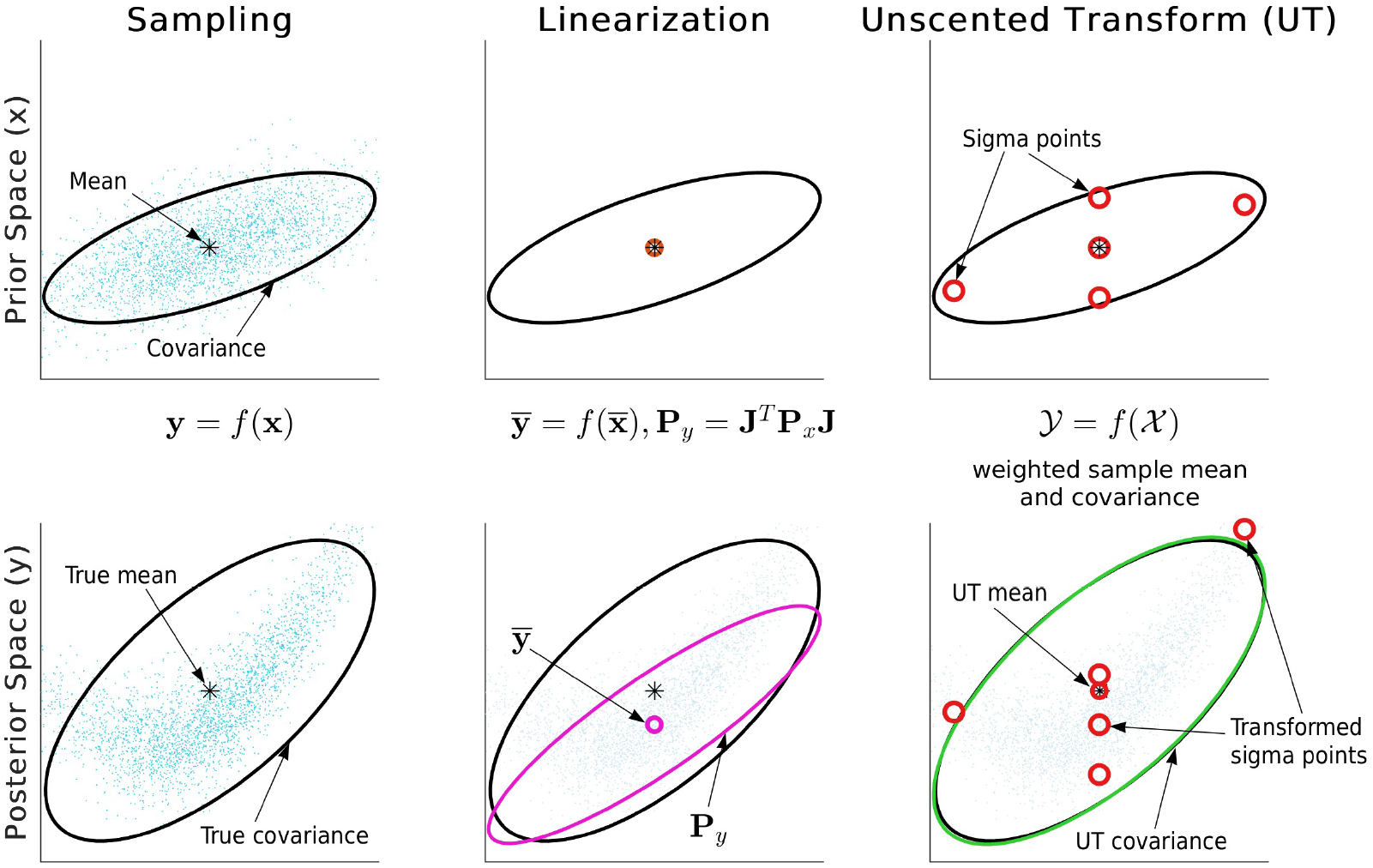
Conceptual comparison of different nonlinear filtering approaches in terms of mean and covariance propagation. (a) Monte Carlo approximation of the true distribution using a large number of samples; (b) first-order linearization used in the EKF, which fails to capture the curvature of the nonlinear transformation; (c) the unscented transformation (UT) used in the UKF, which approximates the distribution using a small set of deterministically chosen sigma points.

To formally define the UT, consider a random vector **x** = (*x*_1_, …, *x*_*L*_). The sigma points form a set of vectors ***ξ*** = [***ξ***_0_, …, ***ξ***_2*L*_] where ***ξ***_*i*_ = (*ξ*_1,*i*_, …, *ξ*_*L,i*_)^*T*^ . These points, along with their corresponding weights 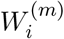 and 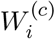, must satisfy the following conditions:

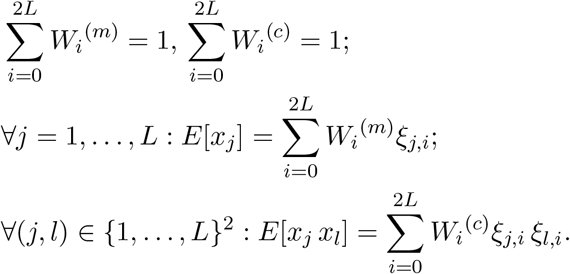

A common selection of sigma points and weights for the random vector **x** is given by:

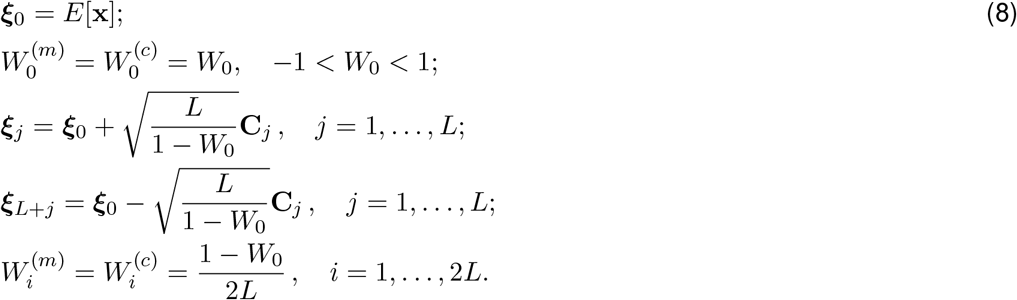

Here, the vector **C**_*j*_ is the *j*-th column of matrix **C**, such that the covariance matrix of **x** satisfies **P**_**x**_ = **CC**^*T*^ . The matrix **C** is typically obtained via the Cholesky decomposition of **P**_**x**_. The central point weight *W*_0_ (set to 0.35 in this work) acts as a tuning parameter that controls the spread of the sigma points around the mean.

Since the state-transition model in this study is linear, the UT is not required during the prediction step. Instead, it is applied only prior to the measurement update step to account for the nonlinearity of the observation model. Consequently, the UKF algorithm employed in this work is formulated as follows:

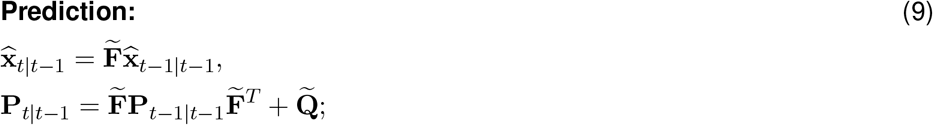

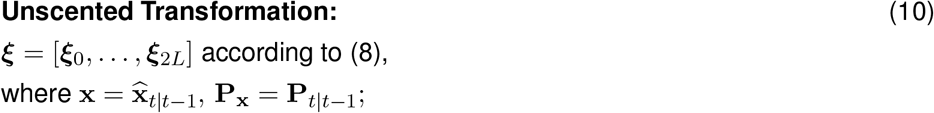

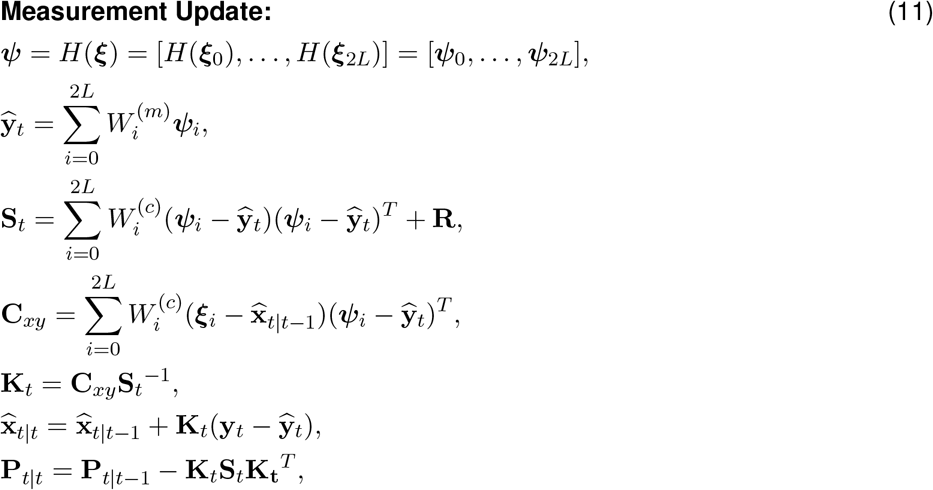

where *H*(·) is the nonlinear observation function defined in Eq. (4).

Thus, the UKF provides the optimal state estimate 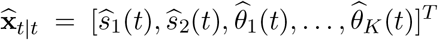 along with the posterior covariance matrix **P**_*t*|*t*_, which quantifies the estimation uncertainty for each latent variable at every time step. Specifically, 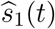 represents the estimated time course of the electrical activity of the dominant brain rhythm source. The auxiliary component 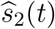, together with 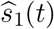allows for the calculation of the instantaneous phase 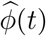 and the envelope 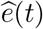 of the rhythm:

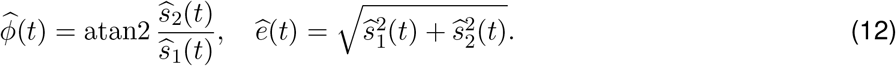

The variables 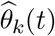 enable the reconstruction of the source position over time, thereby revealing the trajectory of the cortical wave. To reconstruct this trajectory, we utilized a forward model **G**_3_ with freely oriented dipoles, computed in Brainstorm [40] using the overlapping spheres method [41]. Accordingly, for each cortical vertex *r*, we define a local leadfield matrix:

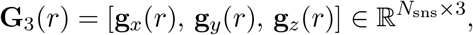

where **g**_*x*_(*r*), **g**_*y*_(*r*) and **g**_*z*_(*r*) are the topographies of a unit dipole placed at vertex *r* and oriented along the *x*-, *y*- and *z*-axes, respectively. At each time point *t*, our goal is to identify the cortical location *r*^∗^ that maximizes the subspace correlation between the model subspace spanned by **G** (*r*) and the estimated topography 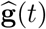, reconstructed from the spatial coefficients:

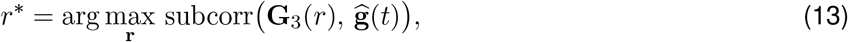

where

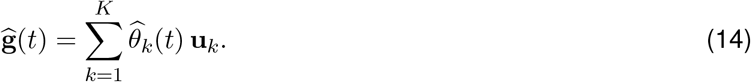

In other words, at each time step *t*, we select the vertex *r*^∗^ that minimizes the principal angle between the column subspace of **G**_3_(*r*^∗^) and the vector 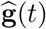. Implementation details for computing subcorr efficiently on large leadfield matrices (for example, ≈ 250, 000 vertices) are provided in Appendix, section Subspace Correlation.

To validate the proposed method, we applied it to both realistically simulated and real MEG/EEG data. The following section describes the datasets used in this study. Unless stated otherwise, the proposed method and the majority of data processing steps, including the data simulation, were implemented in MATLAB [42].

### 2.5 Data

This section provides an overview of the data used in this study, including both simulated MEG data and real MEG/EEG recordings. The simulation data are presented in two scenarios: a wave pattern and two static coherent dipoles.

#### 2.5.1 Wave Pattern Simulation

Realistic MEG data were generated by defining a wave trajectory on the cortical mesh and subsequently modulating the electrical activation — corresponding to the alpha rhythm — along this path. For clarity, the proposed simulation algorithm can be conceptually divided into three stages.

##### Wave Path Generation

In the first stage, an anatomical region of interest (ROI) was selected on a high-density cortical mesh comprising approximately 245,500 vertices. This mesh, generated in Brainstorm based on magnetic resonance imaging (MRI) data, was designated as the area for subsequent modeling. For example, the *calcarine sulcus* with 1,328 vertices served as the representative region, as it has been widely recognized as one of the primary sources of alpha rhythm generation in the brain. The anatomical atlas provided by Brainstorm enabled the precise localization and selection of this region (Figure 3a).

**Figure 3.**
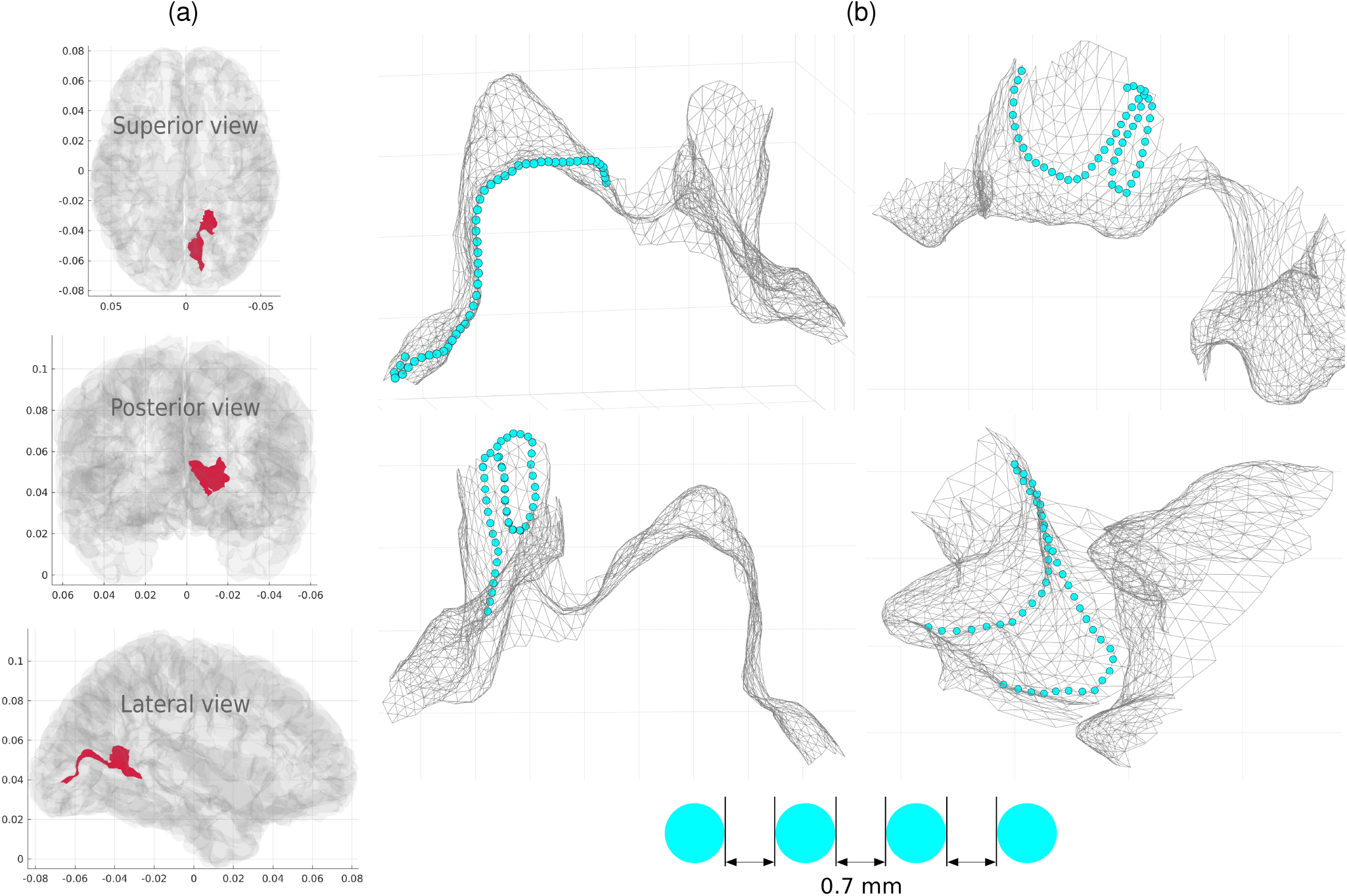
Wave path generation on the cortical mesh. **a)** Localization of the right *calcarine sulcus* (red) relative to the entire cortical surface. **b)** Several variations of the simulated wave path. The distance between consecutive spatial points along each path is 0.7 mm.

The path consists of a one-dimensional sequence of connected vertices with equidistant neighbors. It is characterized by its total length, the number of vertices, and a geometry that conforms to the underlying mesh topology. The path was constructed as follows.

An initial vertex **v**_*s*_ was selected randomly. Next, the average normal vector for this starting vertex and its *N* neighbors was computed as

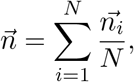

where 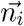 is the normal vector of the *i*-th neighboring vertex. The matrix 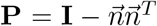 projects any vector onto the tangent plane defined by the normal 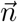.

Subsequently, one of the neighboring vertices, **v**_*n*_, was randomly selected to define the propagation direction. This direction was then determined by the unit vector

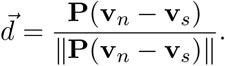

To ensure the path remains as “straight” as possible across the highly convoluted cortical surface, the algorithm defines a virtual plane passing through the starting vertex **v**_*s*_ and oriented along the vector 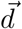. At each subsequent step, the algorithm iteratively selects the neighboring vertex that lies closest to this plane, effectively minimizing lateral deviation from the initial trajectory. A similar approach for path construction was previously employed by Kuznetsova et al. [43].

Finally, the constructed path was resampled to obtain *N*_s_ = 60 points distributed at uniform intervals of mm. To maintain this constant spacing, selected points were interpolated along the edges of the mesh and consequently did not necessarily coincide with the original vertex locations. Figure 3b illustrates several variations of the wave path, which are consistent with patterns reported in the literature [44].

##### Wave Activity Generation

The second stage involved simulating oscillatory electrical activity and defining the spatiotemporal activation matrix along the constructed path. As described in Section 2.3, the alpha rhythm can be modeled using equation (5). For the simulations, we used the following parameters: *c*_**F**_ = 0.99998, *f*_*c*_ = 10, **w**(*t*) ∼ *N* (**0**, 0.1^2^ **I**). This yielded two time series, **z**(*t*) = [*z*_1_(*t*), *z*_2_(*t*)]^*T*^, generated at a sampling frequency of 1000 Hz.

To obtain a realistic TW pattern, the simulation incorporates both 1) a consistent spatial phase shift along the trajectory and 2) a time-varying spatial envelope. This envelope accounts for the sequential recruitment of neighboring neural populations and their subsequent entry into a refractory period, ensuring that only a segment of the path remains active at any given moment. To realize this, the temporal alpha oscillator was augmented with both a distance-dependent phase shift and a spatial envelope along the path. Consequently, while all points on the trajectory share the same underlying oscillatory source, their phases are systematically offset as a function of their position, effectively modeling the propagation of the wave. The spatial envelope of electrical activity along the path was modeled with a Gaussian profile, whose peak position varies smoothly over time. This envelope serves two primary purposes: first, it represents an active cortical patch that extends beyond a single vertex, yielding a more realistic model of neural activity than a single dipole; second, it smooths spatial irregularities in dipole orientations and vertex spacing, which are often present in constrained forward models.

Specifically, to implement the distance-dependent phase shift described above, we define the spatiotemporal alpha-band activation *ψ*(*r, t*) as follows:

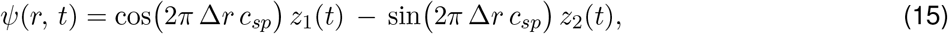

where *r* denotes the spatial coordinate and Δ*r* is the distance (in meters) from the initial vertex to the current location along the path. The phase shift is controlled by the constant *c*_*sp*_, which we chose to yield a physiologically plausible spatial scale of the wave. For our simulations, we set 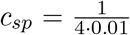 such that a quarter of a wavelength spans 0.01 m. Since the inter-point spacing along the path is constant (0.7 mm), this construction results in a linear phase ramp along the direction of propagation.

The focal activation, representing the active cortical patch, is mathematically expressed by the Gaussian spatial envelope *γ*(*r, t*):

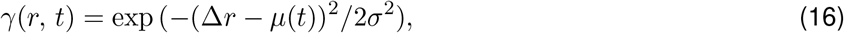

where *σ* is the spatial dispersion parameter and *µ*(*t*) denotes the peak position. The center of the envelope propagates at a constant speed *µ*(*t*) = *v t*, following a back-and-forth trajectory: once the Gaussian center reaches an endpoint of the path, it reverses direction and continues toward the opposite endpoint. We set the propagation speed to *v* = 0.05 m/s in the main simulations; performance for *v* ranging from 0.01 to 0.5 m/s is reported in Table 2. It is worth noting that the actual path traversed by the amplitude peak is shorter than the full predefined path. This restriction prevents the tails of the Gaussian envelope from being truncated at the boundaries. Consequently, out of the total *N*_s_ = 60 vertices, the peak is observed at approximately 36 distinct locations.

The final complex spatiotemporal activation pattern on the cortical mesh is defined by the product *φ*(*r, t*) = *ψ*(*r, t*) · *γ*(*r, t*). For computational implementation, this wave pattern is represented in matrix form as 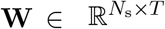, where *N*_s_ is the number of spatial points and *T* is the number of time points. Each column of **W** contains the instantaneous activation amplitudes across the entire path.

The overall generation scheme and the resulting source-level dynamics are visually summarized in Figure 4. Specifically, panel (a) illustrates the modulation of the phase-shifted alpha oscillation by the traveling Gaussian envelope. The evolution of this rhythmic wave pattern across the spatial trajectory is depicted in panels (b) and (c), while panel (d) demonstrates the direct correspondence between the 2D amplitude profile and its 3D anatomical representation on the cortex.

**Figure 4.**
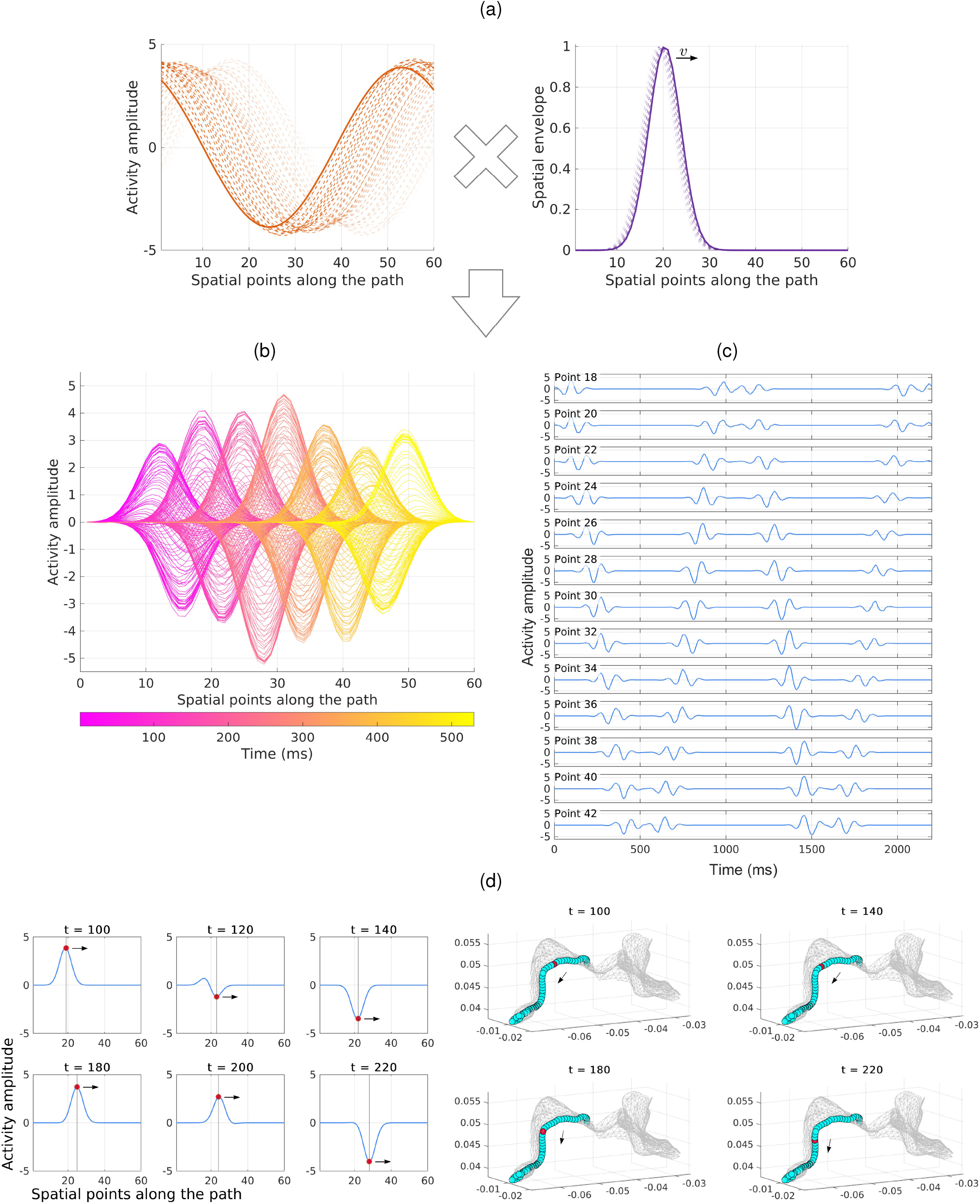
Spatiotemporal activation of the simulated traveling wave at the source level. **a)** Schematic illustration of the activation pattern generation across 60 spatial points via the product *φ*(*r, t*) = *ψ*(*r, t*) · *γ*(*r, t*). The alpha-band oscillatory activity with a distance-dependent phase shift *ψ*(*r, t*) (left, Eq. (15)) is modulated by a spatial envelope *γ*(*r, t*) modeled as a smoothly traveling Gaussian profile (right, Eq. (16)). **b)** The spatiotemporal simulated activity *φ*(*r, t*) plotted across all spatial points along the path. The amplitude varies dynamically over time and space, with color encoding time to illustrate a single traversal along the path over a duration of approximately 530 ms. **c)** Individual time courses of the spatiotemporal activation pattern *φ*(*r, t*) extracted from a subset of selected spatial points along the path, demonstrating the sequential propagation of rhythmic oscillations as the wave passes through distinct cortical locations. **d)** Visualization of the simulated source-level wave progression at selected time points, where the red dot marks the peak location and the arrow indicates the direction of propagation. (Left) The sequence of 2D instantaneous amplitude profiles of the wave along the predefined path. (Right) The corresponding sequence of 3D representations of the exact same simulated state, demonstrating the physical anatomical location of this active peak on the cortical surface.

##### Mapping to Sensors

In the third stage of the simulation, cortical activity was projected onto the sensor space using a standard MEG forward model:

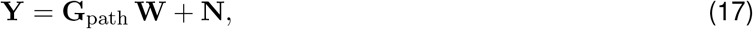

where 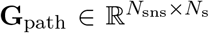 represents the gain matrix corresponding to source locations on the wave path. The columns of **G**_path_ were obtained by interpolating the respective columns of the perturbed leadfield matrix

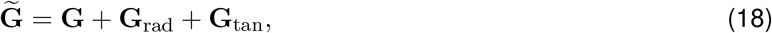

where **G** is the original leadfield matrix computed in Brainstorm via the overlapping spheres method [41] with fixed oriented dipoles. In our simulations, dipole orientations were constrained to be normal to the cortical surface. The stochastic matrices **G**_rad_ and **G**_tan_ represent perturbations along and perpendicular to the leadfield vectors (radial and tangential components, respectively). A detailed derivation of the perturbation modeling is provided in the Appendix (see section Modeling Leadfield Errors). This approach accounts for potential inaccuracies in forward model construction encountered in experimental settings and evaluates the algorithm’s robustness.

Measurement noise **N** was modeled as zero-mean Gaussian noise with a standard deviation (SD) of 0.3 per sensor and time sample. Prior to the addition of noise, the simulated sensor-level signal was normalized to unit variance. Consequently, the SD-based signal-to-noise ratio (SNR) equals SNR = 3.33 in the main simulations; performance for SNR ranging from 0.75 to 4 is reported in Figure 11.

The resulting matrix 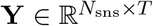 represents the simulated MEG signals. For the purposes of this study, we use Elekta Neuromag gradiometers, resulting in *N*_sns_ = 204. Figure 5 provides an overview of one realization of these simulated sensor-level data. Specifically, panel (b) illustrates the corresponding time series from a subset of occipital MEG sensors. Panel (a) displays a sequence of dynamic MEG topographies reflecting the moving cortical source. Panel (c) shows the singular value decomposition (SVD) spectrum of this data, where three significant components clearly dominate. Consequently, we set the number of eigen-topographies to *K* = 3, a choice that also satisfies the criterion of retaining at least 95% of the total data variance. Finally, panel (d) presents the power spectral density (PSD), confirming the presence of a dominant alpha-band peak around 10 Hz.

**Figure 5.**
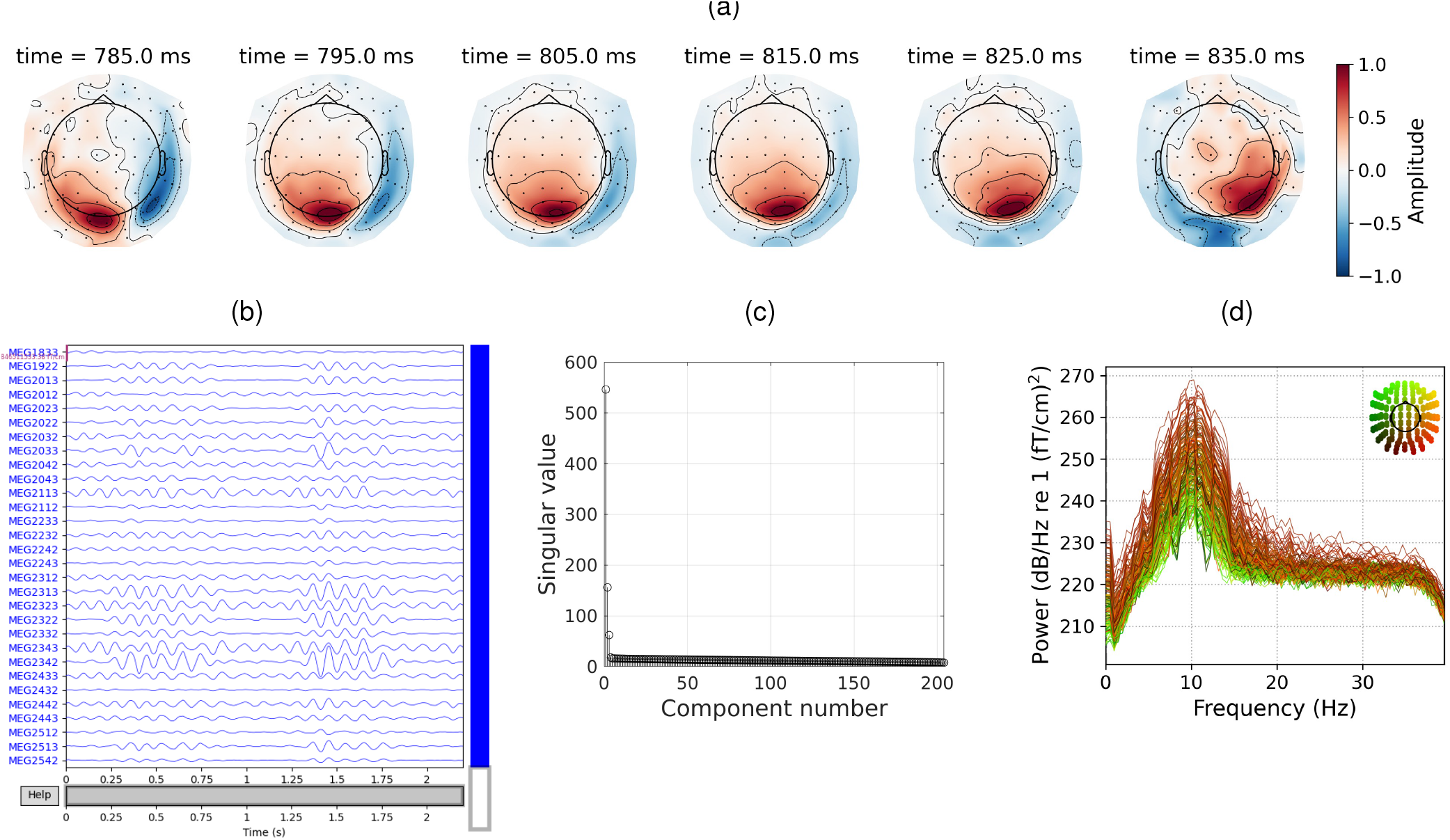
Representation of the simulated MEG data. **a)** Sequence of simulated MEG topographies illustrating the dynamic spatial evolution of alpha-rhythm source activity (10 ms between frames). **b)** Example of the simulated MEG time series from a subset of occipital gradiometers, showing clear alpha spindles. **c)** SVD spectrum of the simulated data. The steep drop-off after the third component motivates the selection of *K* = 3 eigen-topographies. **d)** PSD of the simulated signals, confirming the prominent spectral peak at the 10 Hz carrier frequency.

For subsequent validation, we define the following quantities as ground-truth in the simulation study:

1. Electrical component. For the simulated data, the ground-truth electrical signal is obtained by summing the spatiotemporal activation along the path:

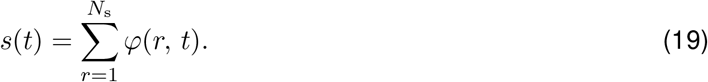
2. Instantaneous phase. Within the simulation procedure, the ground-truth phase is given by

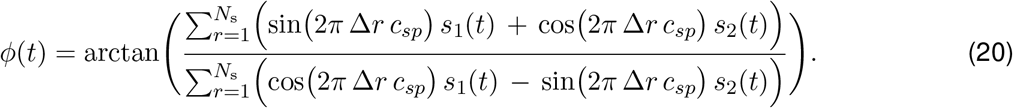
3. Spatial component. The Ground-truth time series of the spatial coefficients ***θ***(*t*) = [*θ*_1_(*t*), …, *θ*_*K*_(*t*)] can be computed as

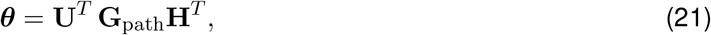

where 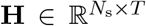 is the matrix form of *H*(*r, t*). The matrix 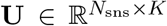 consists of the first *K* left singular vectors obtained from the SVD of the analyzed simulated data.
4. Propagation trajectory. The ground-truth trajectory is defined by the coordinates of the moving Gaussian center, i.e., the peak location of the spatial envelope.

#### 2.5.2 Two Static Coherent Sources Simulation

As highlighted in Section 1, distinguishing a TW from a pair of coherent (phase-locked) static dipoles presents a major challenge for traditional source localization techniques. In this section, we present a simulation designed to explore whether our approach can resolve this specific ambiguity.

The cortical mesh and forward model matrix 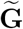 from the expression (18) were identical to those used in the traveling wave simulation. To simulate a scenario where two stationary sources mimic a propagating wave, two discrete dipoles were placed at the starting and ending vertices of the predefined path used in the traveling wave simulation (see Figure 6b). Let **g**_*r*_ and **g**_*l*_ denote the columns of the forward model matrix that correspond to the starting (right) and ending (left) vertices, respectively.

**Figure 6.**
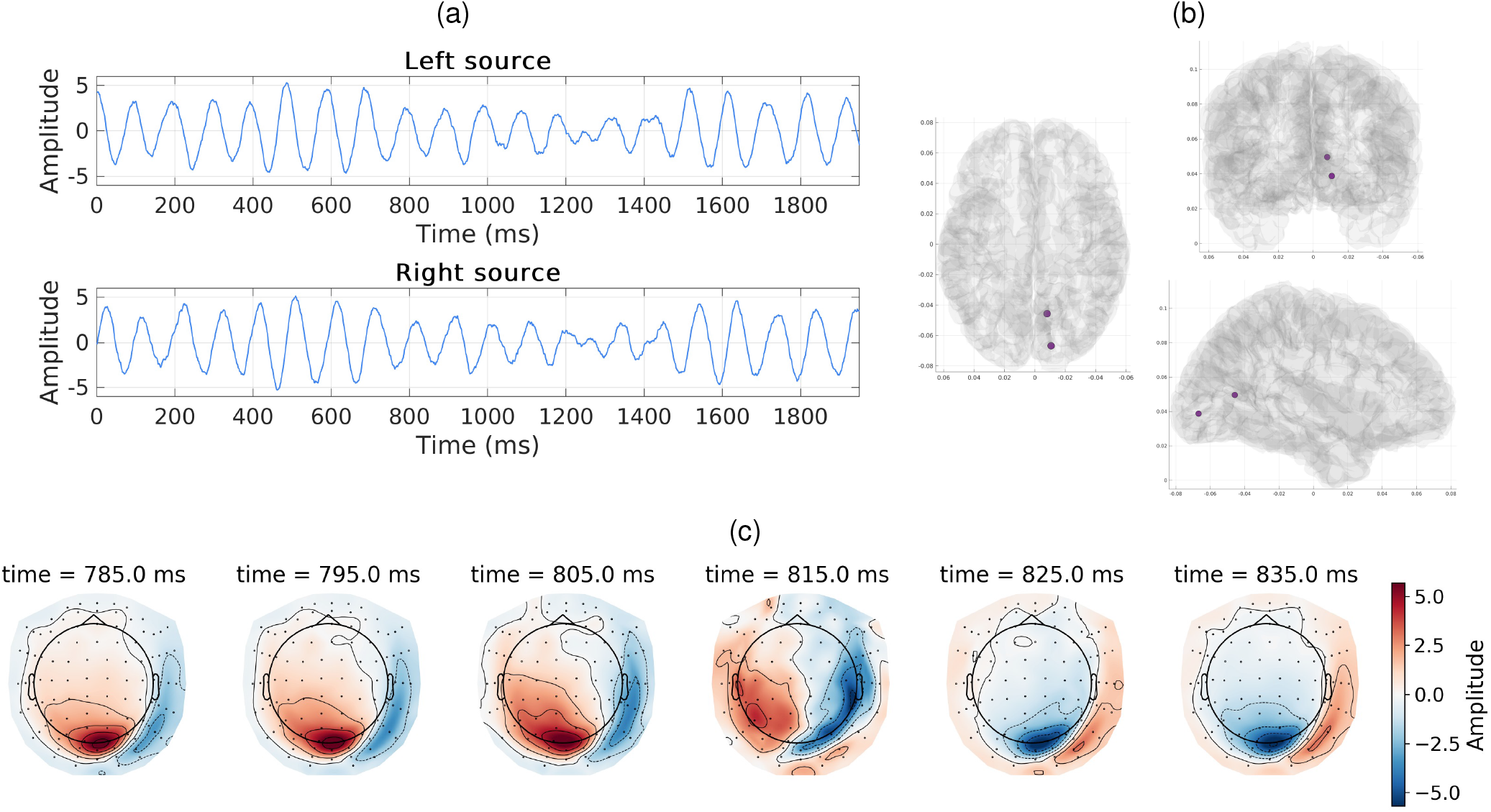
Representation of the simulated data: two coherent static sources. **a)** Time series of alpha oscillations for two static sources. The left source has a phase shift of *π/*2 relative to the right source. **b)** True positions of the static sources on the cortical mesh. **c)** Sequence of MEG topographies illustrating the dynamics of this simulation (10 ms between frames).

The electrical activity time series for the sources were generated using Eq. (5) with the parameters *c*_**F**_ = 0.99998, *f*_*c*_ = 10, and process noise **w**(*t*) ∼ *N* (**0**, 0.1^2^**I**) at a sampling frequency of 1000 Hz. Recall that the state vector for a single oscillator in Eq. (5) is defined as **s**(*t*) = [*s*_1_(*t*), *s*_2_(*t*)]^*T*^, representing the real and imaginary parts of an analytic signal [29]. To introduce a precise *π/*2 phase shift between the two sources, we assigned the real component *s*_1_(*t*) to the left source (*s*_*l*_(*t*) = *s*_1_(*t*)) and the imaginary component *s*_2_(*t*) to the right source (*s*_*r*_(*t*) = *s*_2_(*t*)), as illustrated in Figure 6a.

Similar to the cortical wave simulation, the source activity was mapped onto the sensor space. Rewriting Eq. (17) for two discrete sources yields:

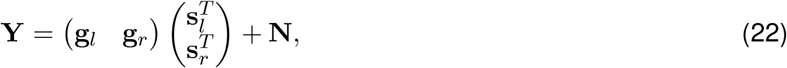

where 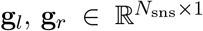, and **s**_*l*_, **s**_*r*_ ∈ ℝ^*T ×*1^. For this simulation, we restricted the primary analysis to the *N*_sns_ = 204 planar gradiometers. The matrix **N** denotes additive measurement noise, with each temporal column sampled from an independent Gaussian distribution, **N**_*t*_ ∼ *N* (**0**, 0.001^2^ **I**) for *t* = 1, …, *T* .

At the sensor level, this activation induces a clear propagating pattern. Specifically, the sequence of topographical distributions (Figure 6c) illustrates the signal’s apparent propagation from one region to another across the parieto-occipital sensors, effectively mimicking the spatial phase gradient typical of a TW. For visual clarity, these topographic maps were generated using only the 102 magnetometers.

In this application, the number of eigen-topographies was set to *K* = 2 as the corresponding set of singular vectors **u**_*k*_ accounts for at least 95% of the variance in this simulation data.

#### 2.5.3 MEG Data

In addition to the simulated data, we also analyzed alpha waves in real EEG and MEG recordings. Specifically, we examined CTF MEG data recorded during the eyes-closed resting state from the NIMH Healthy Research Volunteer Dataset [35].

We used data from a single participant. The recordings were acquired with a 272-channel CTF MEG system (CTF MEG, Coquitlam, BC, Canada) using third-order gradient balancing. MEG signals were sampled at 1200 Hz with a quarter-Nyquist filter at 300 Hz. Standard preprocessing was conducted using MNE-Python software [45, 46], including bandpass filtering between 2 and 40 Hz. Biological artifacts, such as cardiac and ocular activity, were removed via independent component analysis (ICA, [47]), as implemented in MNE-Python, with five artifact components excluded.

In this study, our primary focus is on the alpha rhythm. To identify alpha spindles, we used the implementation of the Spectro-Spatial Decomposition (SSD, [48]) algorithm available in MNE-Python. The targeted frequency band was set to 9–12 Hz, with surrounding frequencies at 8 and 13 Hz. Subsequently, alpha spindles were identified through visual inspection of the SSD components. For further analysis, we extracted multiple time series segments from the full MEG recording, selecting intervals where alpha spindles were present. We also selected a ROI comprising *N*_sns_ = 85 MEG channels covering the parieto-occipital region (see Figure 7b). All of the channels in the CTF MEG system are axial gradiometers.

**Figure 7.**
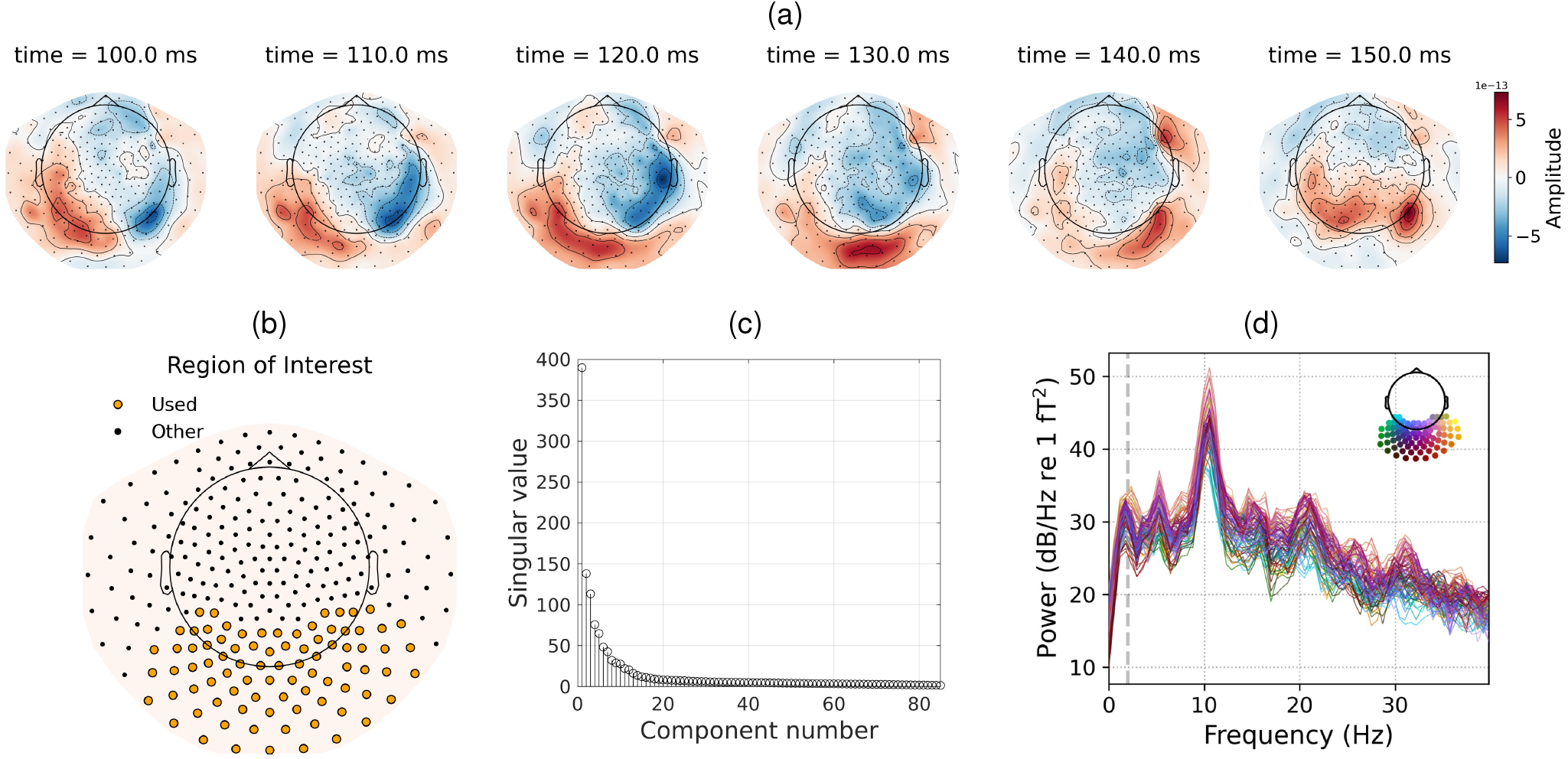
Representation of the real MEG data. **a)** Sequence of MEG topographies illustrating the dynamic evolution of alpha-rhythm source activity (10 ms between frames). **b)** Region of interest (ROI) comprising 85 MEG channels covering the parieto-occipital area. **c)** SVD spectrum of the selected segment of real MEG data (ROI). It shows that the most prominent variance is concentrated in the first three components, motivating the selection of *K* = 3 eigen-topographies. **d)** PSD of the MEG data segment confirmed the presence of a dominant alpha-band peak around 11 Hz.

Furthermore, the participant’s individual structural MRI (T1-weighted) from this dataset was used to compute a forward head model. First, cortical and head surfaces were extracted from the MRI using FreeSurfer software (version v7.4.0, [49]). Then, the head model was generated in Brainstorm using the overlapping spheres method [41] with free dipole orientations. The number of vertices is approximately 245,500, covering the cortical surface.

In this application, the number of eigen-topographies was set to *K* = 3 as the corresponding set of singular vectors **u**_*k*_ accounts for at least 95% of the variance in the real MEG data (see Figure 7c).

The selected data segment, with a duration of 2.024 seconds, is characterized by a distinct peak around 10 Hz (Figure 7d) and exhibits a characteristic sequence of source topographies in the occipital region (Figure 7a).

#### 2.5.4 EEG Data

Eyes-closed resting-state EEG data were acquired from a single participant using a 38-channel montage at a sampling rate of 1000 Hz. Electrodes were arranged according to the extended 10-20 system, with increased sensor density over the occipital region (see Figure 8b for details). The signals were originally recorded using a monopolar montage and were subsequently transformed using common average referencing. The recording was performed via an NVX52 amplifier with NeoRec application software (Medical Computer Systems Ltd, Moscow, Russia), maintaining electrode impedances below 15 kΩ throughout the session.

**Figure 8.**
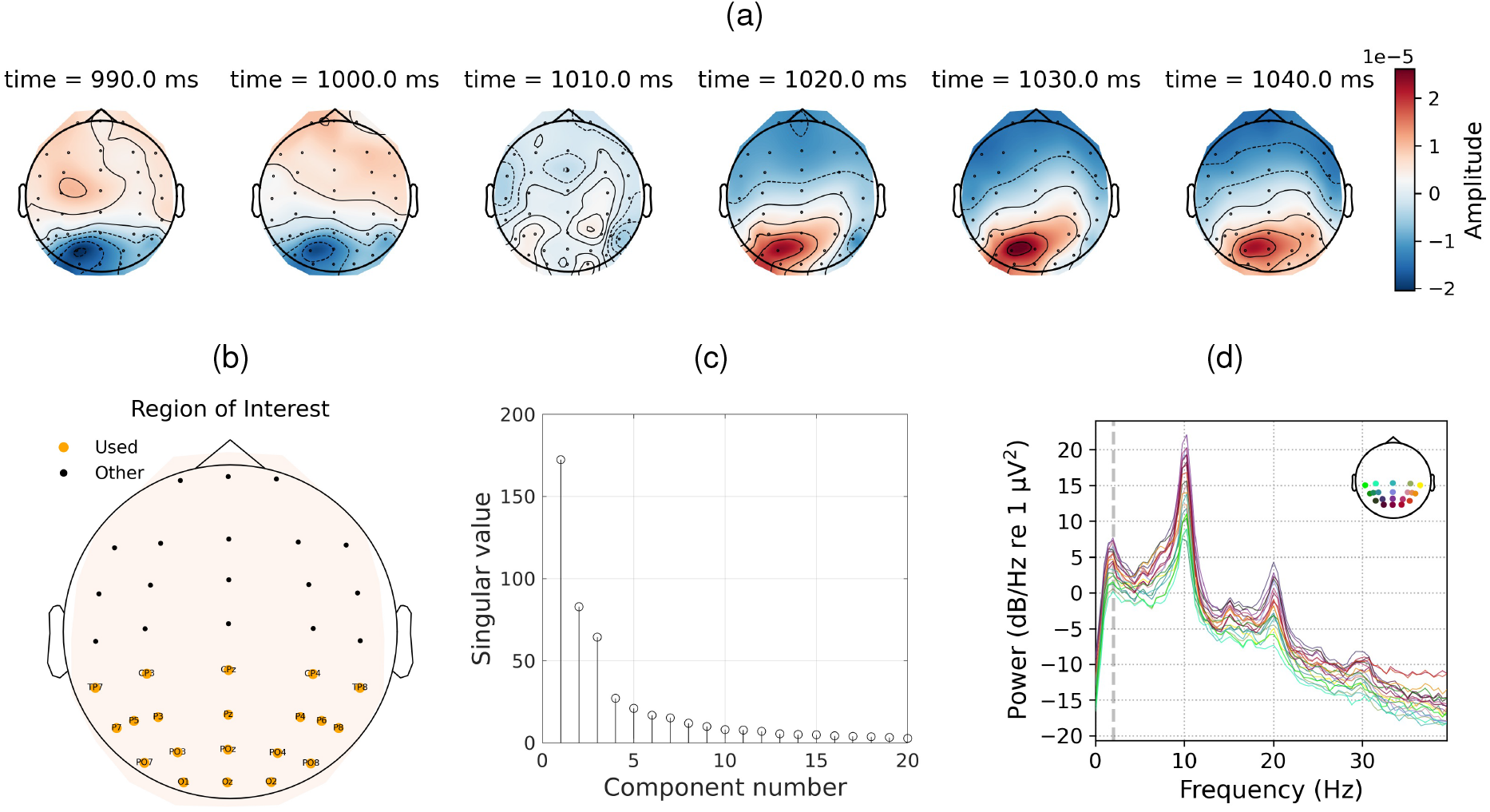
Representation of the real EEG data. **a)** Sequence of EEG topographies illustrating the dynamic evolution of alpha rhythm source activity (10 ms between frames). **b)** Region of interest (ROI) comprising 20 EEG channels covering the parieto-occipital area. **c)** SVD spectrum of the selected segment from the alpha condition (ROI). It shows that the most prominent variance is concentrated in the first three components, motivating the selection of *K* = 3 eigen-topographies. **d)** PSD of the EEG data segment confirmed the presence of a dominant alpha-band peak around 10 Hz.

Preprocessing was conducted using MNE-Python. The data were bandpass-filtered between 2 and 40 Hz and notch-filtered at 50 Hz. Subsequently, biological artifact correction was performed via ICA, resulting in the exclusion of five components related to eye movements and cardiac activity. For the core analysis, we selected an ROI comprising *N*_sns_ = 20 EEG channels covering the parieto-occipital region (Figure 8b).

A forward head model was computed based on the participant’s individual structural MRI (T1-weighted, acquired on a Philips Intera 1.5T scanner). First, cortical and head surfaces were extracted from the MRI using FreeSurfer software. Subsequently, the head model was computed in Brainstorm using the OpenMEEG BEM method [50, 51] with free dipole orientations. The resulting cortical mesh comprised approximately 270,000 vertices.

In this application, the number of eigen-topographies was set to *K* = 3 as the corresponding set of singular vectors **u**_*k*_ accounts for at least 95% of the total variance in the real EEG data (see Figure 8c).

From the continuous recording, we selected a segment with a pronounced alpha rhythm characterized by a non-stationary topography in the parieto-occipital region. The chosen data segment exhibits a distinct spectral peak around 10 Hz (Figure 8d) and displays a characteristic sequence of source topographies across the parietooccipital sensors (Figure 8a).

### 2.6 Parameter Setting

In this study, the implementation of the UKF models requires the estimation of several key parameters: the central frequency of the rhythm *f*_*c*_; the measurement noise covariance matrix **R**; the process noise covariance matrices **Q** and **B**, which correspond to the spatial coefficients and electrical component, respectively; and two scalar parameters, *c*_**A**_ and *c*_**F**_, associated with the state transition matrices **A** and **F**.

Within the recursive Bayesian inference framework, the Kalman filter acts as a generative model describing the probabilistic process that yields the sequence of observations **Y** = (**y**_1_, …, **y**_*T*_). Because the latent state variables are not directly observed, the model parameters are estimated by maximizing logarithm of the marginal likelihood, which measures the probability of the observed data given the model:

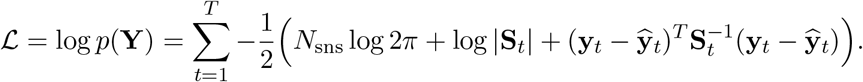

Here, **S**_*t*_ denotes the innovation covariance matrix at time step *t*, while 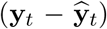 represents the prediction residuals (innovations). By optimizing the parameters to maximize ℒ, the model is tuned to best explain the observed data.

To solve this optimization problem, we employed a two-stage computational approach implemented in MATLAB, combining a grid search technique with the interior-point method [52]. The grid search was initially utilized to establish reasonable physiological boundaries and identify a stable starting point for the algorithm. Based on this preliminary analysis, the search space for the subsequent interior-point optimization was strictly constrained. The noise covariance matrices **R, Q**, and **B** were assumed to be diagonal, with their elements bounded within [10^−3^, 1], [10^−3^, 1], and [10^−6^, 10^−2^], respectively. The scaling parameters *c*_**A**_ and *c*_**F**_ were restricted to the range [1 − 10^−2^, 1 − 10^−8^] to ensure the stability and smoothness of the state transitions. The central frequency *f*_*c*_ was defined based on the frequency corresponding to the dominant peak in the PSD of the analyzed data segment.

### 2.7 Baseline Methods for Comparison

In order to assess the effectiveness of the proposed method and evaluate the improvement in tracking accuracy, we selected classical linear Kalman filtering (KF), Minimum Norm Estimation (MNE), and traditional dipole fitting for comparative analysis.

#### Linear Kalman Filter

The classical KF focuses solely on the electrical dynamics, ignoring the local movements of the source. This standard approach effectively serves as an ablation study, allowing us to demonstrate the necessity of incorporating dynamic spatial topography. In this restricted setting, instead of the nonlinear observation model (Eq. (4)), we employed the following linear model:

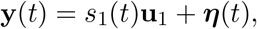

where **u**_1_ is the first singular vector of the data matrix, and it corresponds to the scenario with *K* = 1. This formulation does not allow for tracking spatial components and implicitly assumes that the source is static. Equation (5) still serves as the process model. Although this yields estimates for both 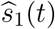 and 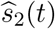, only 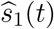) was used for the comparative analysis.

#### Minimum Norm Estimation

The MNE approach with Tikhonov regularization [36] solves the matrix form of the observation equation:

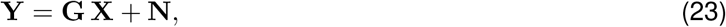

where 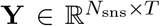 is the data matrix, 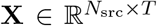 is the matrix of unknown neuronal source activity, 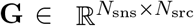 is the forward model matrix, and **N** is the noise matrix. For the distributed inverse problem, the number of sources *N*_src_ equals the number of vertices (e.g., 1328 for the calcarine sulcus). The inverse solution for this underdetermined system is given by:

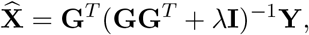

where the regularization parameter *λ >* 0 was selected using the L-curve method [53]. The mean 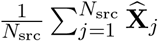 was used as the estimate of the source amplitude for the following comparative analysis.

#### SVD-Filtered Dipole Fitting

To evaluate localization quality, we applied a dipole fitting approach enhanced with spatial filtering. The analyzed data segment was first band-pass filtered in the 8–12 Hz range, resulting in the signal 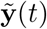. To reduce noise and ensure a fair comparison, the filtered data was projected onto the first *K* = 3 left singular vectors **U**_1:*K*_, thereby achieving spatial filtering analogous to the basis utilized in our proposed algorithm. The cortical surface was scanned at each time step to find the active source. The location was estimated as the vertex that maximized the subspace correlation (as defined in Eq. (13)) between the forward model subspace spanned by **G**_3_(*r*) and the projected measurements 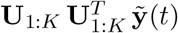.

It is important to note that the baseline localization methods (RAP-MUSIC and dipole fitting) were applied to the full sensor array to maintain the stability of the spatial covariance estimates and ensure global coverage of the forward model. In contrast, our proposed framework was constrained to a spatially restricted subset of channels (ROI-focused, Figure 7b and Figure 8b). This methodological distinction is intentional: while traditional methods are fundamentally designed to scan the global cortical space based on broad spatial covariance, our approach relies on a generative model specifically tuned for local spatiotemporal tracking. By focusing on a region of interest, we enhance sensitivity to local oscillatory dynamics, whereas traditional global-scanning methods require the full sensor topology to prevent unrealistic source placement.

## 3 Results

### 3.1 Simulation Studies

#### 3.1.1 Wave Pattern Tracking

To evaluate the performance of our method, we first tested the time-series reconstruction accuracy. We ran 1000 simulations with random realizations of process and measurement noise, as well as varying simulated propagation paths along the calcarine sulcus (see Figure 3b for examples). Over these runs, the proposed algorithm (hereafter labeled as UKF-Inv in all figures and tables) accurately estimated the electrical component of the simulated propagating cortical activity, achieving a median correlation of 0.94 between the true and estimated time series. This indicates consistently high performance and robustness across trials. Figure 9a illustrates a specific example of this reconstruction, comparing the estimated electrical amplitude 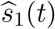 with the groundtruth source activity *s*(*t*) (Eq. (19)). For this particular instance, the time series are highly correlated (corr = 0.98), highlighting the potential of the proposed method to accurately track the electrical component in real data. Furthermore, the estimated phase closely matches the ground-truth phase of the source time series. As presented in Figure 9e, the difference between the true phase *ϕ*(*t*) (Eq. (20)) and its estimate 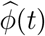 (Eq. (12)) is minimal, demonstrating the method’s ability to reliably capture dynamic phase variations in the signal. This capability is particularly relevant for studies involving phase-based connectivity and oscillatory dynamics.

**Figure 9.**
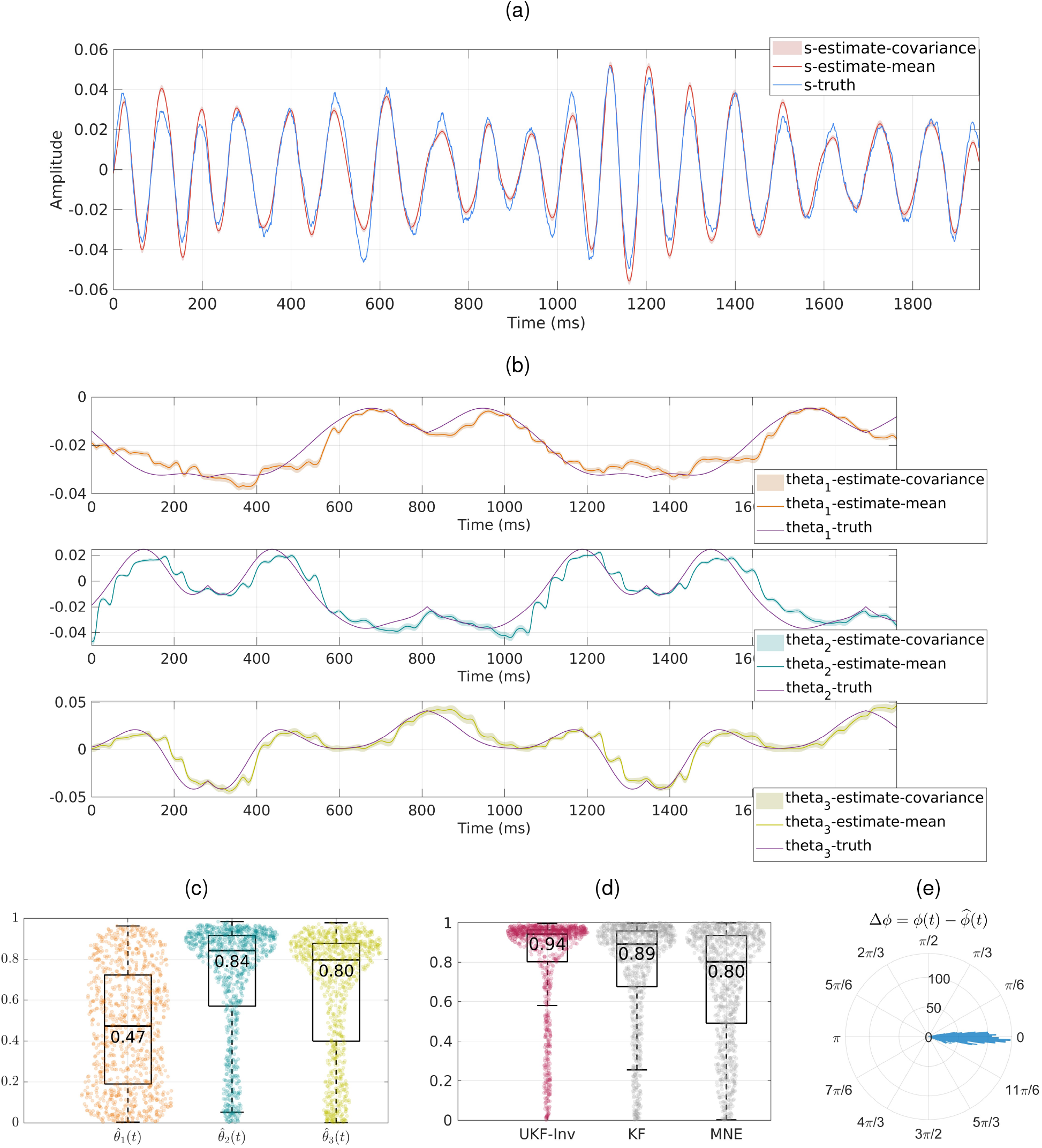
Results of time series estimation for simulated data. **a)** Estimation of the amplitude of the electrical activity *ŝ*_1_(*t*), where *s-truth* denotes the ground-truth alpha oscillations of the source, and *s-estimate* represents the estimate obtained using our UKF-based method. **b)** Estimation of the spatial coefficients 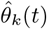, where *theta-truth* denotes the ground-truth time series and *theta-estimate* is the estimate obtained using our UKF-based method, shown top to bottom for each *k* = 1, 2, 3. **c)** Distribution of the spatial correlation coefficients for the estimated parameters 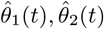, and 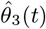 using the UKF across 1000 iterations. Boxplots denote the medians and interquartile ranges, while overlaid swarm plots illustrate the individual data points. The median values are annotated above each corresponding distribution. **d)** Comparison of correlation coefficients for the electrical signal component *ŝ*_1_(*t*) across three estimation methods over 1000 iterations. Boxplots and overlaid swarm plots illustrate the distribution of correlations between the time series of the true and estimated electrical components. Evaluated methods include: MNE — minimum norm estimation with Tikhonov regularization; KF — linear Kalman filter without spatial component tracking; and UKF-Inv — the proposed algorithm. Median values are annotated for each distribution. **e)** Polar histogram of the phase estimation error. The sharp peak centered at 0 radians demonstrates the high phase-tracking precision of the UKF-Inv framework.

Next, we evaluated the method’s potential to capture the traveling wave trajectory by reconstructing the time series of spatial coefficients. As detailed in the Methods section, we retained *K* = 3 eigen-topographies. Figure 9c illustrates the performance of the UKF in estimating these variables across 1000 iterations. The results indicate a varying degree of tracking accuracy among the three coefficients. Specifically, the estimates for 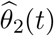 and 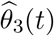 demonstrate strong spatial correlations, achieving median values of 0.84 and 0.80, respectively. Conversely, the estimation of 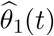 yields a lower median correlation of 0.47. Empirically, this first component is frequently observed to capture the core spatial topography, typically exhibiting a more stable and less temporally variable amplitude dynamic. Consequently, due to this lower temporal variance, the Pearson correlation metric becomes inherently more sensitive to minor background noise fluctuations. In contrast, the second and third components carry the highly dynamic, sign-alternating variations required to capture the physical displacement of the wave. Since these estimated coefficients are utilized to approximate the evolving topography (Eq. (14)) — which is subsequently used for source localization via subspace correlation (Eq. (13)) — the robust tracking of these dynamics is particularly vital.

An illustrative example of the estimated spatial coefficients in relation to ground-truth values is shown in Figure 9b. Notably, the coefficients exhibit periodic fluctuations consistent with the source’s back-and-forth motion along the predefined path. For this specific trial, all components are tracked with high precision, yielding correlation values of 0.94, 0.93, and 0.93, respectively. Overall, the obtained results indicate that the proposed method effectively recovers the essential spatiotemporal variability underlying the propagating source, thereby ensuring accurate subsequent localization.

To assess the accuracy of the reconstructed path, localization was performed using two stages: restricting the search to a predefined ROI (e.g., around the calcarine sulcus) and conducting a global search over the entire cortical surface. Across 1000 simulation runs, the median time-averaged geodesic error on the cortical mesh between the ground-truth coordinates of the moving Gaussian center and the localized vertex was 3.02 mm for the ROI-restricted search and 3.15 mm for the global search, with medians of 24.5 and 26 detected unique points, respectively. Considering that the cortical mesh is highly non-uniform — with inter-vertex distances ranging from 0.036 to 8.345 mm (averaging approximately 1 mm) — and that the ground-truth path was constructed via interpolation with 0.7 mm spacing between points, these results indicate both high localization accuracy and robustness to scale.

Additional visual evidence of this scalability and precision is provided in Figure 10. Note that the simulated source performed a continuous back-and-forth movement. For visualization purposes, this continuous dynamic is separated into two segments: an upward trajectory on panel (a) and a downward trajectory on panels (b) and (c). Furthermore, to evaluate the robustness of the proposed method, the reconstruction was performed under two different spatial constraints. While panels (a) and (b) demonstrate the results on a local mesh of the calcarine sulcus (approximately 1,300 vertices), panel (c) confirms the method’s efficiency and accuracy when scaling up to the full cortical surface mesh (approximately 245,500 vertices). For this illustrative example, the mean geodesic error equals 1.5 mm for both the ROI-restricted and the global searches, each detecting 46 trajectory unique points. Ultimately, utilizing dynamic spatial coefficients yields a smoother trajectory estimate, and allowing free dipole orientations in the forward model further improves localization accuracy.

**Figure 10.**
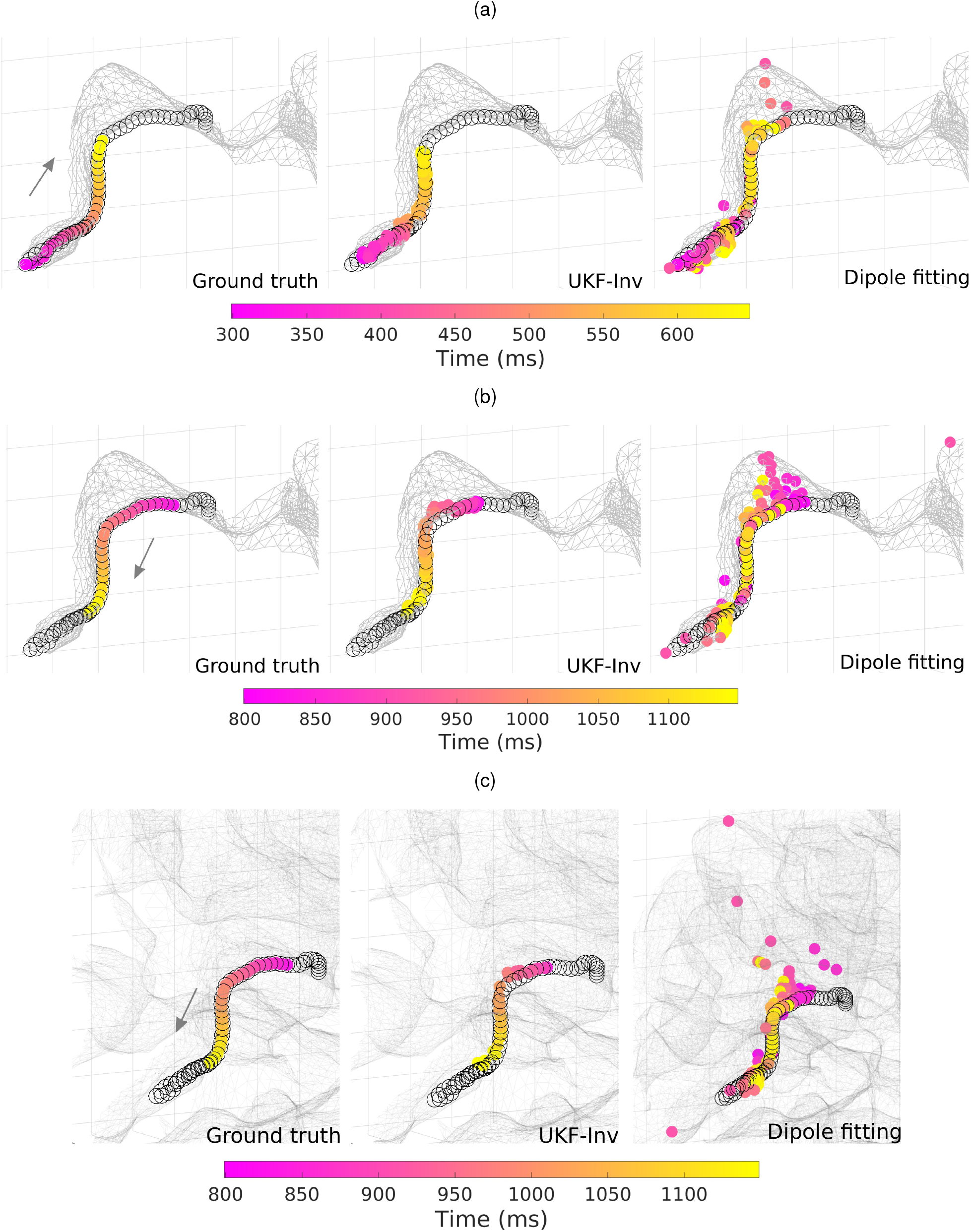
Results of trajectory reconstruction for simulated MEG data. Two segments of source propagation along a cortical path are presented: an upward trajectory **(a)**, and a downward trajectory **(b, c)**. Panels **(a)** and **(b)** display the localization results computed on a local cortical mesh restricted to the calcarine sulcus, whereas panel **(c)** illustrates the downward trajectory evaluated on the full cortical surface mesh. In each panel, the left column displays the ground truth trajectory of the alpha source, where each point indicates the true location of the spatial envelope peak. The center column illustrates the reconstructed trajectory using the proposed UKF-Inv method, while the right column represents the localization via the dipole fitting approach. Across all plots, point colors encode time in milliseconds, visualizing the spatiotemporal dynamics of the source.

In addition to the stages described above performed on the right calcarine sulcus, the entire pipeline was also applied to the cortical surface of the left central sulcus with approximately 3,300 vertices. The highly efficient performance of the algorithm in this region demonstrates its capability to track not only occipital alpha rhythms but also sensorimotor mu rhythms and other propagating patterns. The resulting metrics were essentially indistinguishable from those reported above and are summarized in Table 1, further supporting the robustness and versatility of the proposed method.

**Table 1:** Comparison of source localization performance between the proposed UKF-Inv framework and traditional dipole fitting. Results are evaluated for sources simulated in two distinct cortical regions, utilizing both local ROI-restricted and global search spaces.

|  | UKF-Inv |  | Dipole fitting |  |
| --- | --- | --- | --- | --- |
|  | Calcarine | Central | Calcarine | Central |
| Local search | 3.02 mm | 3.18 mm | 5.09 mm | 5.34 mm |
| Global search | 3.15 mm | 3.81 mm | 6.48 mm | 6.71 mm |

(b) Median number of unique vertices detected during source localization procedure.
|  | UKF-Inv |  | Dipole fitting |  |
| --- | --- | --- | --- | --- |
|  | Calcarine | Central | Calcarine | Central |
| Local search | 24.05 | 36.5 | 103 | 134 |
| Global search | 26.00 | 36.5 | 129 | 155 |

To evaluate the robustness of the proposed framework across varying propagation speeds of the spatial envelope — modeled as a smoothly traveling Gaussian profile in Section 2.5.1 — we tested its performance on simulated data. The experimental scenario replicated the steps previously described in Section 2.5.1 across seven distinct speeds, ranging from 0.01 m/s to 0.5 m/s. For each speed, 30 independent 1-second simulations were generated.

Table 2 summarizes the evaluation metrics across all tested propagation speeds. Specifically, we report: (i) the correlation between the estimated electrical component *ŝ*_1_(*t*) and the ground-truth envelope amplitude of the rhythm; (ii) the correlations between the estimated spatial coefficients 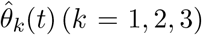 and their corresponding ground-truth values; and (iii) the source localization error, quantified as the time-averaged geodesic distance between the localized vertex and the ground-truth source position.

**Table 2:** Comprehensive performance metrics of the proposed framework across varying propagation speeds *v* of spatial envelope (16). Reported metrics include the correlation between the estimated electrical component *ŝ*_1_ and the ground-truth rhythm amplitude, correlations between the estimated spatial coefficients 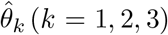 and their true values, and the localization error — time-averaged geodesic distance from the estimated vertex to the ground-truth location. Values represent the median (Q1–Q3) computed over 30 simulation runs in the calcarine sulcus.

|  | Propagation Speed of Spatial Envelope (m/s) |  |  |  |  |  |  |
| --- | --- | --- | --- | --- | --- | --- | --- |
|  | 0.01 | 0.03 | 0.05 | 0.07 | 0.1 | 0.3 | 0.5 |
| Electrical corr., $\hat{s}_1$ | 0.96<br>(0.91–0.98) | 0.95<br>(0.92–0.97) | 0.94<br>(0.91–0.96) | 0.95<br>(0.93–0.97) | 0.97<br>(0.85–0.98) | 0.94<br>(0.64–0.97) | 0.91<br>(0.59–0.95) |
| Spatial corr., $\hat{\theta}_1$ | 0.68<br>(0.38–0.88) | 0.56<br>(0.23–0.76) | 0.67<br>(0.45–0.87) | 0.61<br>(0.26–0.83) | 0.59<br>(0.33–0.82) | 0.32<br>(0.14–0.63) | 0.05<br>(0.03–0.18) |
| Spatial corr., $\hat{\theta}_2$ | 0.96<br>(0.86–0.98) | 0.87<br>(0.67–0.93) | 0.87<br>(0.64–0.92) | 0.85<br>(0.48–0.92) | 0.90<br>(0.45–0.95) | 0.64<br>(0.26–0.83) | 0.34<br>(0.09–0.59) |
| Spatial corr., $\hat{\theta}_3$ | 0.87<br>(0.66–0.95) | 0.86<br>(0.58–0.94) | 0.80<br>(0.60–0.89) | 0.78<br>(0.45–0.85) | 0.83<br>(0.33–0.93) | 0.51<br>(0.13–0.76) | 0.25<br>(0.08–0.44) |
| Localization error | 1.9 mm<br>(1.3–2.5) | 2.5 mm<br>(2.1–3.3) | 2.8 mm<br>(2.3–3.7) | 2.7 mm<br>(2.4–4.0) | 2.8 mm<br>(2.1–3.8) | 3.4 mm<br>(2.6–4.4) | 4.4 mm<br>(3.6–5.8) |

Based on these results, we observe that the algorithm maintains robust performance at slower speeds of the spatial envelope. However, at higher velocities, such as 0.3 and 0.5 m/s, the evaluation metrics begin to decline. This behavior follows directly from the timescale-separation assumption on which the model rests. Within a single carrier cycle the envelope peak advances by *v/f*_*c*_, which equals 5 mm at *v* = 0.05 m/s but 30 and 50 mm at 0.3 and 0.5 m/s — comparable to or exceeding the 42 mm length of the simulated path. At these speeds the spatial and the electrical components vary on the same timescale and can no longer be decoupled. The degradation is not uniform across the estimated quantities: the electrical component remains well recovered throughout (median correlation ≥ 0.91 at every speed tested) and the localization error grows gradually, from 1.9 mm at 0.01 m/s to 4.4 mm at 0.5 m/s, whereas the spatial coefficients — and hence the fidelity of the reconstructed trajectory — are affected most.

To evaluate the effect of measurement noise on the performance of the proposed approach, we tested it across varying SNR levels. Following the simulation scenario described in Section 2.5.1 — where SNR is defined as the inverse of the standard deviation of the additive noise applied to the normalized sensor-level data — we investigated six SNR levels: 4, 3, 2, 1.35, 1, and 0.75. For each level, 30 independent 1-second simulations were generated.

Figure 11 illustrates the evaluation metrics across all tested measurement noise levels. Specifically, panel a describes the effect on the electrical time series reconstruction, while panel b reflects the influence on spatial coefficient tracking. As expected, the correlation metrics for both components tend to decline with decreasing SNR. For the electrical time series, lower SNR levels lead to increased inter-simulation variability of correlation values across individual runs, although the median correlation remains high. For the spatial weights, alongside this growing variability, we observe a marked drop in correlation for the third coefficient *θ*_3_. This reduction occurs because *θ*_3_ corresponds to the third eigen-topography **u**_3_, which is closest to the noise-dominated SVD components, making it highly susceptible to noise corruption.

**Figure 11.**
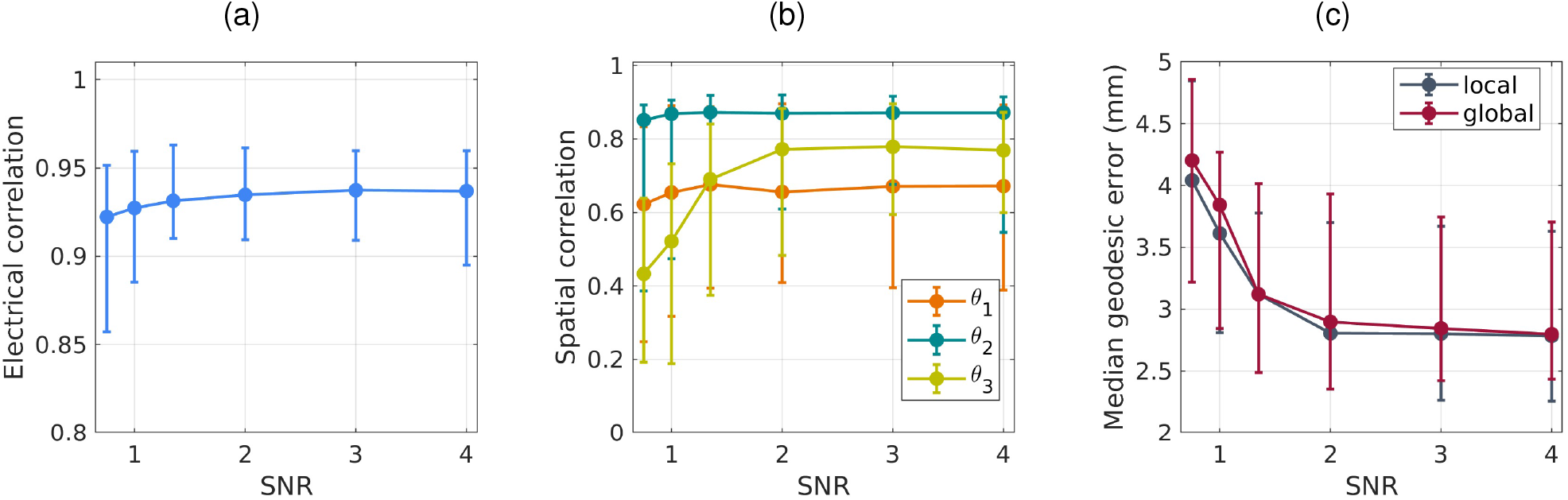
Effect of measurement noise variation. **a)** Median and interquartile range for correlation value for electrical component *ŝ*_1_(*t*). **b)** Medians and interquartile ranges for correlation values for spatial components 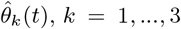. **c)** Medians and interquartile ranges for time-averaged geodesic errors of localization on *local* and *global* meshes, corresponding calcarine sulcus and whole brain. The curves on panels **(a)**-**(c)** were obtained by simulation 30 independent realizations.

Panel c displays the localization error as a function of SNR for both local and global source search spaces, performed on the calcarine sulcus and the whole-brain cortical mesh, respectively. Both localization errors exhibit a synchronous increase as the SNR approaches 1. This behavior is a logical consequence of both the noisy estimation of the spatial coefficients and the noise corruption of the spatial basis itself (specifically, the third eigen-topography **u**_3_). Overall, the results demonstrate the robustness of the algorithm, with a noticeable drop in localization accuracy occurring only when the signal and noise levels become comparable. Nevertheless, even under such low-SNR conditions, global search across the entire cortical mesh recovers a compact trajectory.

#### 3.1.2 Comparison with Baseline Methods

Figure 9d compares the performance of our approach (UKF-Inv) against Minimum Norm Estimation (MNE) and a classical linear Kalman filter (KF) in reconstructing the electrical component of the signal. Notably, the comparison with the classical KF serves as an ablation study to isolate the benefits of explicitly modeling dynamic topography. The distribution of correlation coefficients across 1000 iterations demonstrates that the proposed algorithm yields the most accurate and stable results. Specifically, UKF-Inv achieved a median correlation of 0.942 (IQR: 0.80–0.96), outperforming both the KF ablated model (median 0.89, IQR: 0.67–0.95) and MNE (median 0.80, IQR: 0.50–0.93). The variance visualized by the swarm plots further confirms that the new method not only provides higher accuracy but also exhibits greater robustness across iterations. Ultimately, this confirms that our approach better estimates the electrical activity of cortical sources exhibiting spatial variations — an advantage directly achieved through the more informed measurement and state models.

To evaluate localization quality, we compared the reconstructed trajectories obtained via our dynamic spatiotemporal tracking model and the traditional dipole fitting approach with SVD-filtering (see Section 2.7 for implementation details). Table 1 summarizes the localization comparison results across 1000 simulation runs. In contrast to the robust tracking of the UKF-Inv algorithm, dipole fitting demonstrated a significantly more diffuse trajectory reconstruction. For the calcarine sulcus ROI-restricted search, dipole fitting yielded a median time-averaged geodesic error of 5.09 mm with a median of 103 unique detected vertices, whereas our approach achieved 3.02 mm with 24.5 unique detected vertices. Similar discrepancies were observed for the global search (6.48 mm vs. 3.15 mm) and across the central sulcus conditions, confirming the consistency of this improvement.

Figure 10 provides visual evidence that corroborates these statistical findings. The traditional technique produced a scattered and spatially inconsistent trajectory that failed to adequately capture the smooth propagation pattern characteristic of a cortical wave. In contrast, our algorithm produced a much smoother and more coherent reconstruction of the source trajectory. This improvement is directly attributable to the explicit tracking of the dynamical topography **g**(*t*) and the implicit constraints on its slow temporal variability, which effectively regularize the localization problem in both local and global search contexts.

#### 3.1.3 Two Static Coherent Sources Analysis

To evaluate whether our algorithm can resolve the ambiguity between genuine propagation and stationary coupled sources producing phase-shifted patterns at the sensor level, we applied it to the simulated two-dipole scenario described in Section 2.5.2. Our primary objective was to confirm that the model correctly identifies the static nature of these sources, rather than misinterpreting their phase offset as spatial propagation.

We hypothesized that our single-source model would capture the most prominent of the two sources: its electrical component would be represented by *ŝ*_1_(*t*), while the localized vertices would tightly cluster around the corresponding ground-truth location. The results confirmed this expectation. Figure 12a demonstrates a high correlation (corr = 0.94) between the true electrical activity of the left source and the Kalman filter estimate. Furthermore, panel c shows that all 11 localized vertices are concentrated near the true left source, which is situated closer to the cortical surface and thus dominates the sensor-level signal. Examining the dynamics of the spatial components 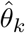 for *k* = 1, 2 (Figure 12b) reveals noticeable local fluctuations, although the overall course remains globally flat. Such fluctuations are expected, as our framework is specifically designed to track a single propagating wave rather than to resolve multiple stationary generators. While this interference slightly broadens the spatial spread of the localized vertices, the globally stationary dynamics of the spatial weights ensure a compact localization cluster that clearly does not resemble a propagating wave.

**Figure 12.**
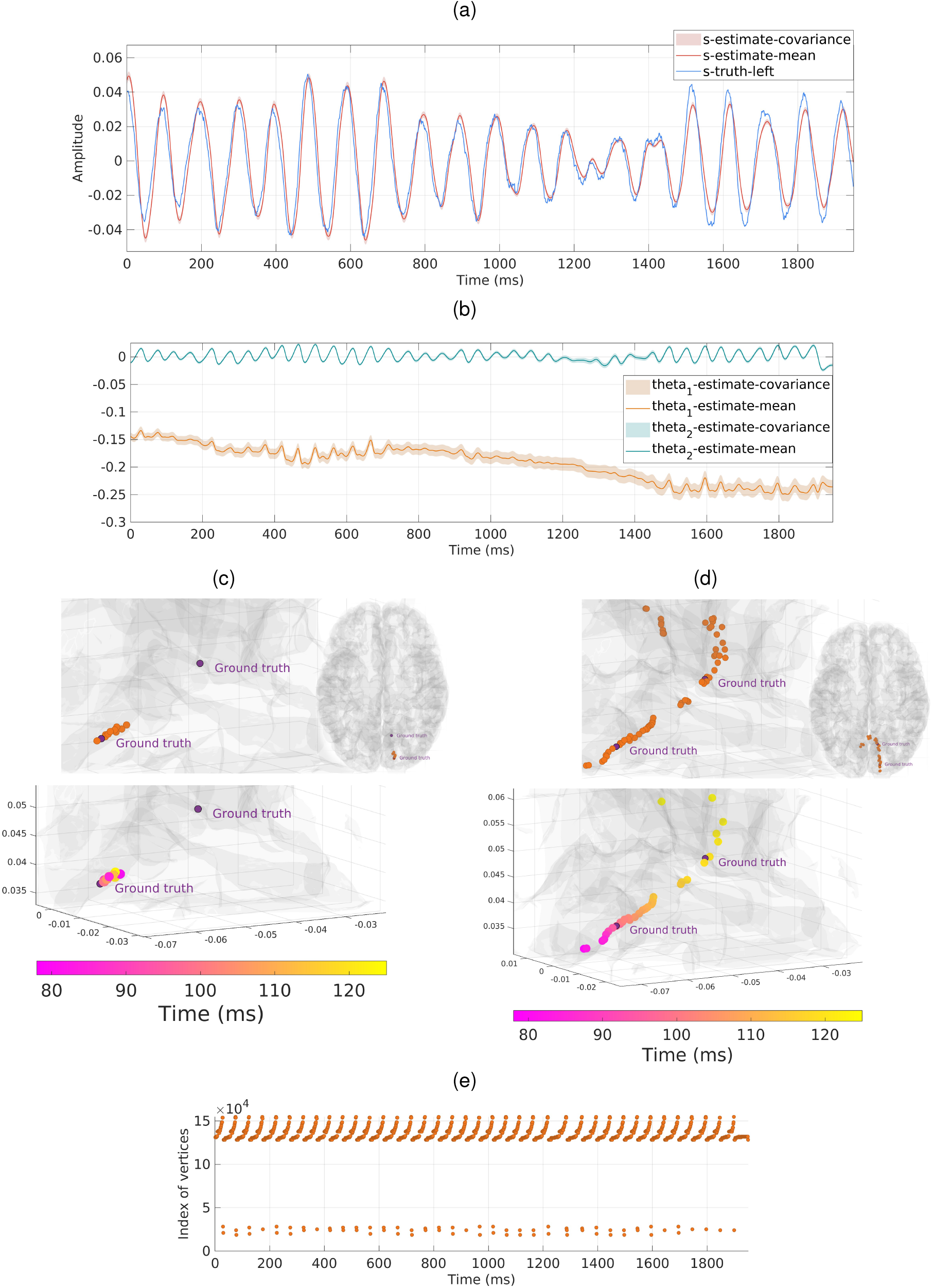
Results of simulated data analysis with two coherent static sources. Panels **a**–**c** illustrate the outcomes obtained using the proposed dynamic spatial tracking framework, while panels **d** and **e** present the results from the independent dipole-fitting approach. **a)** The estimated electrical activity time series (*ŝ*_1_(*t*), *s-estimate*), extracted by the model, plotted alongside the true activity of the left source (*s-truth-left*) for comparison, as it provided the best match. **(b)** Estimated spatial coefficients 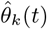 (*k* = 1, 2). **c)** Source localization using the proposed method. Top: the full set of estimated source locations, which are concentrated around the true left source. Bottom: a subset of localization points corresponding to a 50 ms time window. **d)** Source localization using the dipole-fitting approach. Top: the full set of fitted dipole positions. Bottom: a subset of points corresponding to a single cycle (approximately 50 ms). **e)** The localized vertex index plotted as a function of time, demonstrating the cyclic repetition of the fitted dipole position. For spatial plots **c** and **d**, the top panels display the locations in both zoomed-in and superior views. In the bottom panels (zoomed-in view), marker colors encode the temporal progression in milliseconds.

The importance of this disambiguation becomes evident when comparing our results to the traditional dipolefitting approach, where a single equivalent current dipole is estimated at each independent time slice. When applied to data generated by two stationary sources with a constant phase offset, the optimization procedure attempts to explain the mixed topography using only one source. Consequently, an independently fitted dipole will artificially track this shift, creating the illusion of a propagating wave. We observed exactly this artifact when applying dipole fitting to our simulated data (band-pass filtered in the 8–12 Hz range). The algorithm identified 68 distinct vertices spatially distributed between the two true source positions (Figure 12d top). The estimated dipole location exhibited a smooth, cyclic transition across these intermediary vertices every ∼50 ms (panel e), a pattern directly driven by the phase offset imposed in the simulation. This spurious wave-like trajectory is clearly illustrated in panel d (bottom) for one cycle. In empirical data, such a pattern could easily be misinterpreted as a genuine propagating cortical process.

These results demonstrate the robustness of the proposed method even in situations where sensor-level patterns falsely mimic a propagating cortical wave, whereas conventional approaches fall into this trap. This highlights the strong potential of our algorithm for analyzing complex empirical data, where concurrent source configurations often create confounding scenarios.

Given these limitations of traditional static models, we propose using our dynamic spatial tracking framework as an initial diagnostic step to determine whether an observed MEG/EEG pattern stems from a propagating wave or static sources. If the spatial coefficients indicate stationary dynamics, researchers can safely switch to algorithms specifically designed for discrete, potentially correlated sources, such as methods from the MUSIC family. This combined pipeline provides a comprehensive solution: it allows researchers to confidently proceed with our dynamic model to track the spatiotemporal trajectories of genuine waves while relying on established discrete-source algorithms for scenarios identified as static.

### 3.2 Application to Empirical Data

#### 3.2.1 MEG Data Analysis

To confirm the potential of the proposed method in practical scenarios, we applied it to real MEG data. Figure 13 presents the results of this analysis. Panel (a) shows the estimated electrical component time series, which exhibits a center frequency of 10.55 Hz, a characteristic of the alpha rhythm. Panel (b) depicts the estimated spatial coefficients that enable us to track the evolving topography. By combining these variables, we were able to track the propagation path of the source, as illustrated in Figures 14a, b. The observed activity is predominantly concentrated in the right occipital region — an area classically associated with alpha-band generators — and, to a lesser extent, propagates toward the right parietal areas.

**Figure 13.**
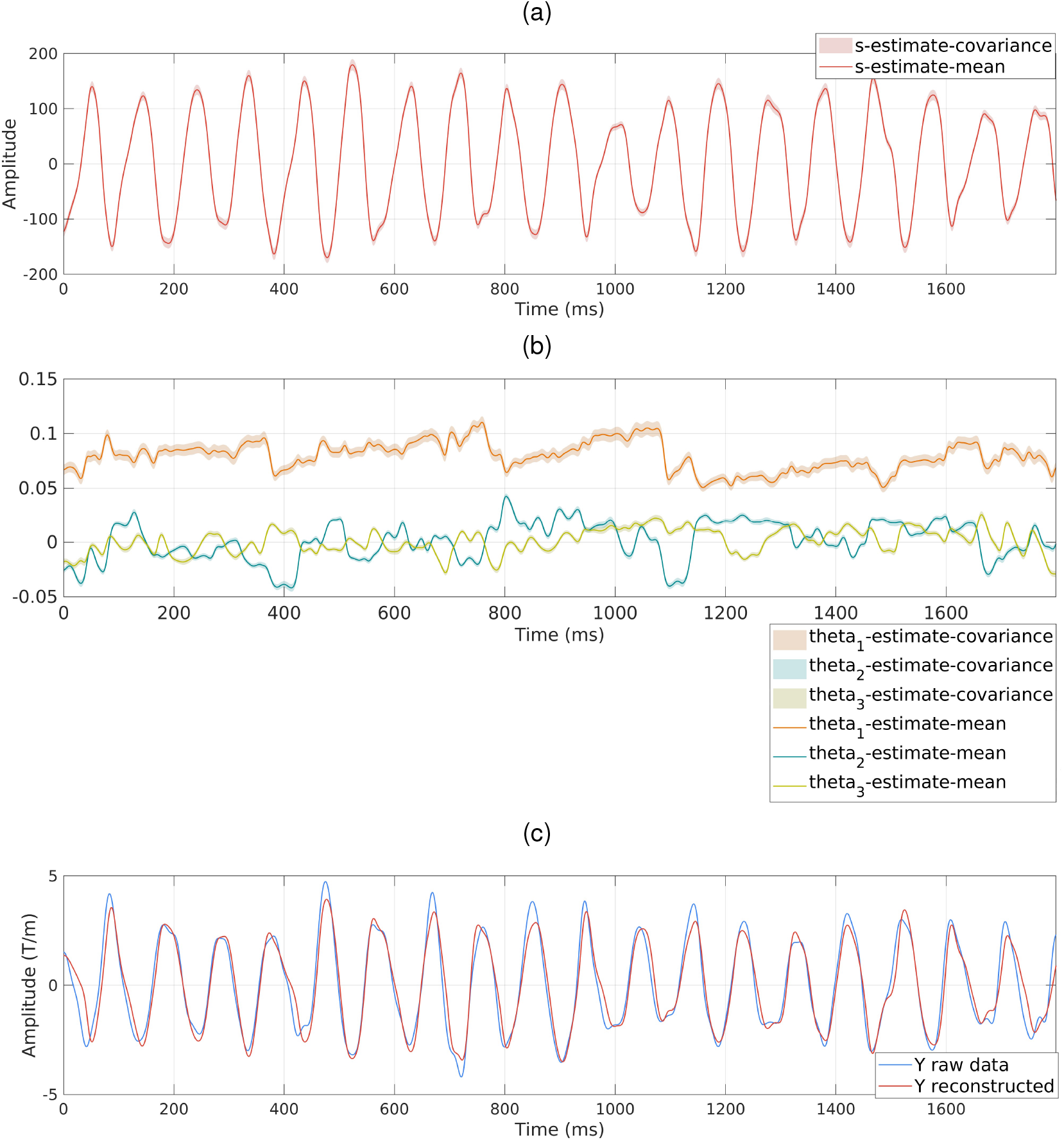
Results of real MEG data analysis obtained with the proposed method. **a)** Estimation of the electrical activity amplitude *ŝ*_1_(*t*) corresponding to the alpha-oscillation source. Here, *s-estimate-mean* and *s-estimate-covariance* denote the mean and covariance of this estimate, obtained using the proposed UKF-based method. **b)** Estimation of the spatial components 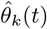, for *k* = 1, 2, 3. Here, *theta*_*k*_*-estimate-mean* and *theta*_*k*_*-estimate-covariance* denote the mean and covariance of these estimates, obtained using the proposed UKF-based method. **c)** Reconstructed time series for a single sensor. Here, *Y raw data* denotes the original time series from that sensor, and *Y reconstructed* represents the reconstructed values obtained using the observation model (4) with the estimates *ŝ*_1_(*t*) and 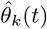, but without the noise term ***η***(*t*).

**Figure 14.**
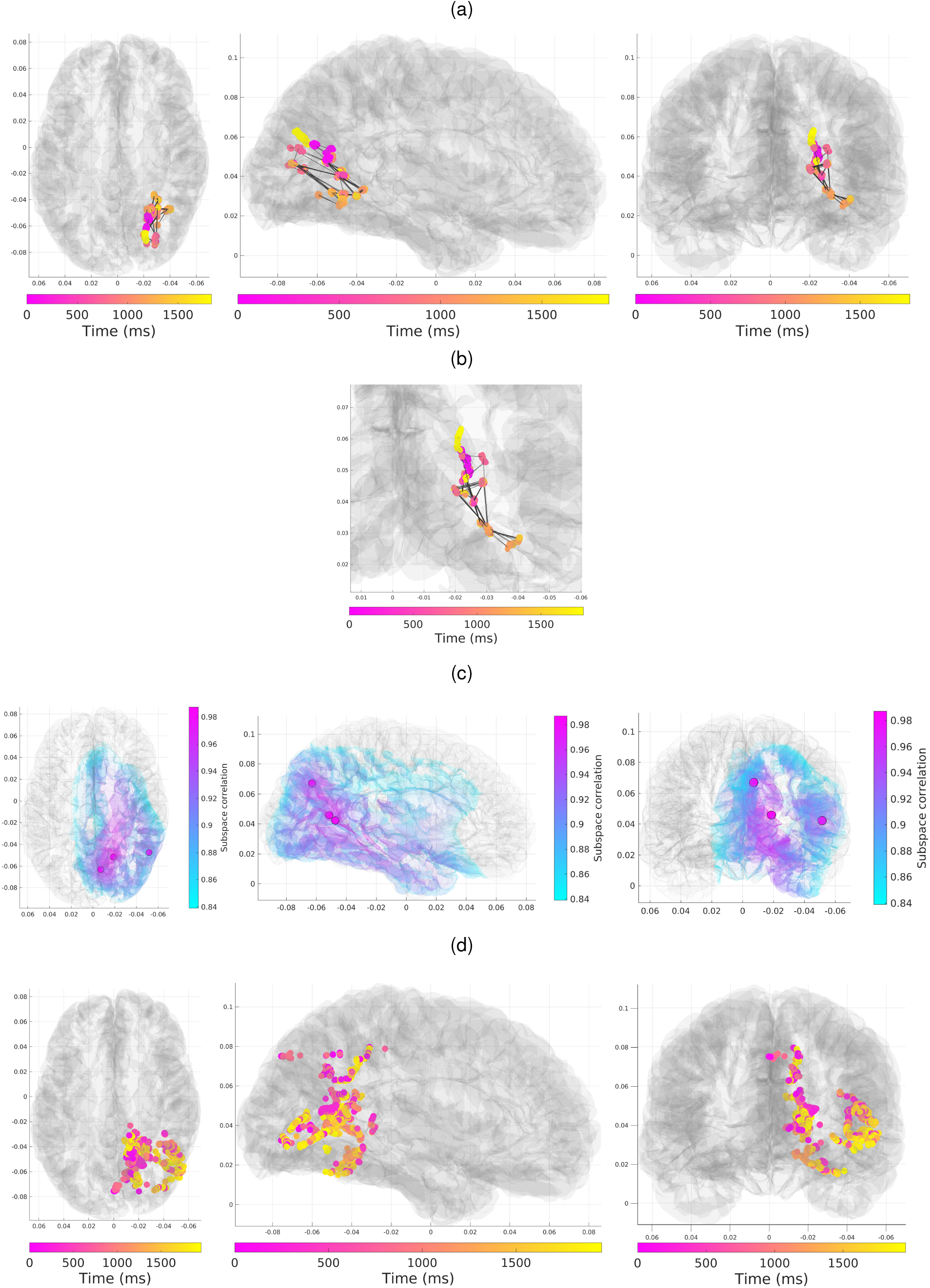
Source localization for real MEG data. **a, b)** Results of the proposed UKF-Inv method. The reconstructed trajectory of the alpha-band source is presented in full-brain views **(a)** and a zoomed-in posterior view **(b)**.Successive locations are connected to visually illustrate the source propagation. **c)** Localization results using the RAP-MUSIC algorithm, which identifies three distinct static dipoles. The reported subspace correlation value corresponds to the first extracted dipole only. **d)** Results of the traditional dipole-fitting approach, where a single equivalent current dipole is estimated at each independent time slice. Across panels **a, c**, and **d**, the cortical mesh is displayed from superior, lateral, and posterior perspectives to provide a comprehensive view. For the spatiotemporal tracking methods (**a, b**, and **d**), each point represents the estimated source location at a specific time instant, with the color encoding the temporal progression in milliseconds.

To validate the anatomically identified active areas, we applied traditional source localization methods, including RAP-MUSIC, to the same data segment. This approach utilized all MEG channels after band-pass filtering in the 8–12 Hz range. As shown in Figure 14c, RAP-MUSIC (using a subspace correlation threshold of 0.85) recovered three distinct static dipoles. The locations of these dipoles are characteristic of alpha-rhythm sources and are anatomically consistent with the core regions identified by our algorithm. While RAP-MUSIC is inherently designed to estimate static equivalent current dipoles, this agreement confirms that our dynamic spatiotemporal tracking operates within biologically plausible and well-established spatial boundaries.

Furthermore, we evaluated the traditional dipole fitting approach enhanced with SVD-filtering on the same sensor data (see Section 2.7 for implementation details). The results, presented in Figure 14d, exhibit a highly scattered localization pattern. While the general spatial cloud remains bounded within the occipital and parietal regions (compatible with alpha-related activity), the traditional dipole fitting yields a highly scattered and spatially fragmented pattern. This stands in contrast to the substantially more compact and spatially structured pattern reconstructed by our approach.

As a final validation step, we assessed the accuracy of the sensor-level data reconstruction. This was achieved by substituting the estimated state variables back into the observation model (4), excluding the noise term. The reconstructed signal is given by:

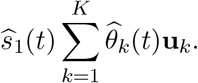

A comparison between the original and reconstructed time series for a single representative sensor is provided in Figure 13c. As evident from the plot, the main temporal dynamics are captured accurately, with the reconstructed signal explaining 80% of the variance (*R*^2^ = 0.80) ^1^ in the sensor data. Notably, in an ablation test where the dynamic spatial coefficients 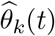 are replaced by their temporal averages, the explained variance drops to 73% (*R*^2^ = 0.73). This reduction explicitly quantifies the substantial contribution of the evolving spatial topography to the overall signal. Regarding the remaining variance in the full model, higher-frequency fluctuations are not reproduced by the Kalman filtering step, which is an expected outcome since the algorithm was purposefully designed to track the dominant alpha-band oscillatory activity and slow spatial transitions rather than rapid, broadband noise fluctuations.

#### 3.2.2 EEG Data Analysis

The analysis of the EEG data followed the same procedure described for the MEG data, including the estimation of the electrical and spatial components, the reconstruction of the source trajectory, and a comparison with baseline localization methods. Figures 15 and 16 summarize the results of this pipeline.

**Figure 15.**
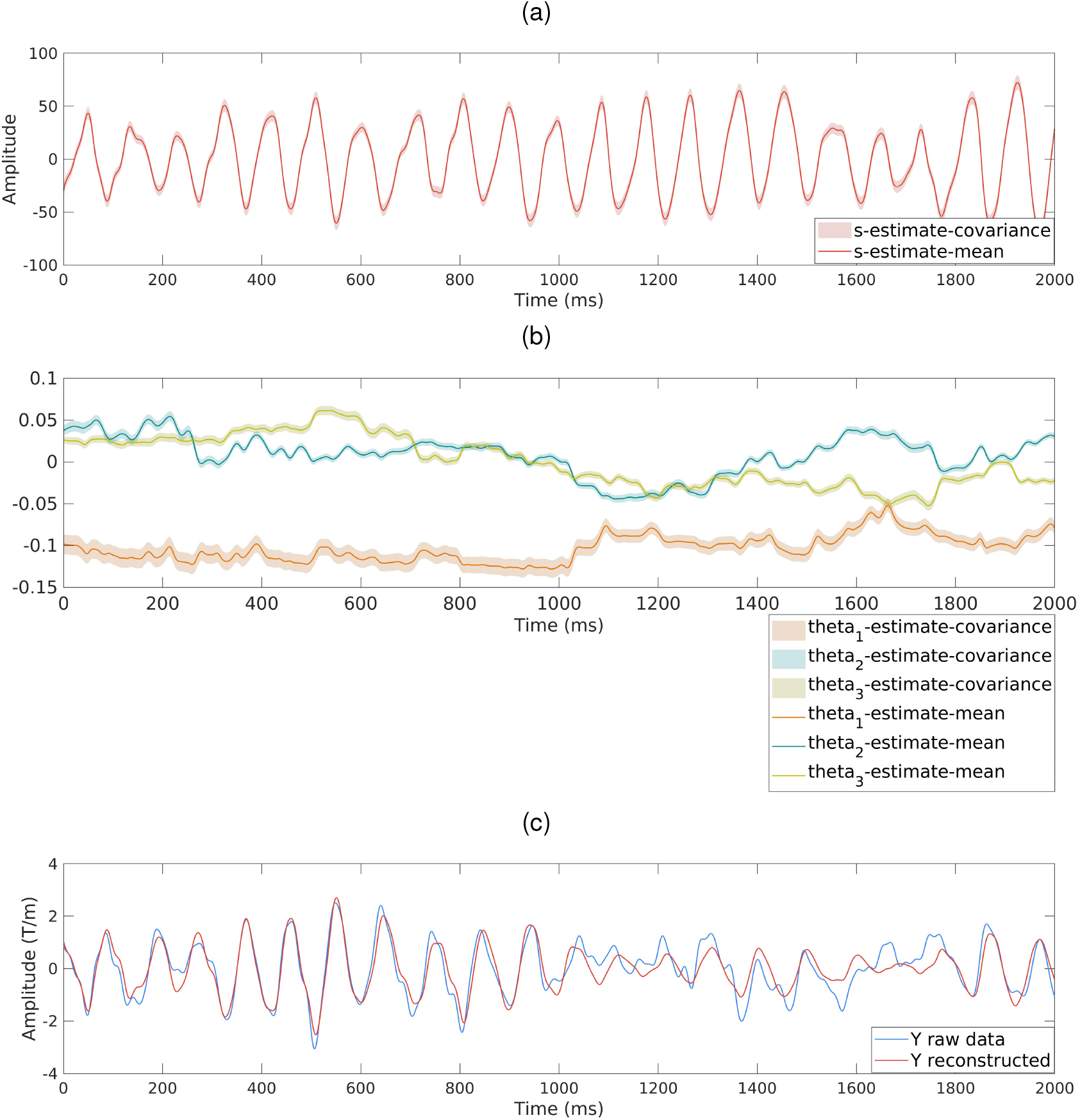
Results of real EEG data analysis obtained with the proposed method. **a)** Estimation of the electrical activity amplitude *ŝ*_1_(*t*) corresponding to the alpha-oscillation source. Here, *s-estimate-mean* and *s-estimate-covariance* denote the mean and covariance of this estimate, obtained using the proposed UKF-based method. **b)** Estimation of the spatial components 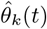, for *k* = 1, 2, 3. Here, *theta*_*k*_*-estimate-mean* and *theta*_*k*_*-estimate-covariance* denote the mean and covariance of these estimates, obtained using the proposed UKF-based method. **c)** Reconstructed time series for a single sensor. Here, *Y raw data* denotes the original time series from that sensor, and *Y reconstructed* represents the reconstructed values obtained using the observation model (4) with the estimates *ŝ*_1_(*t*) and 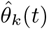 (*t*), but without the noise term ***η***(*t*).

**Figure 16.**
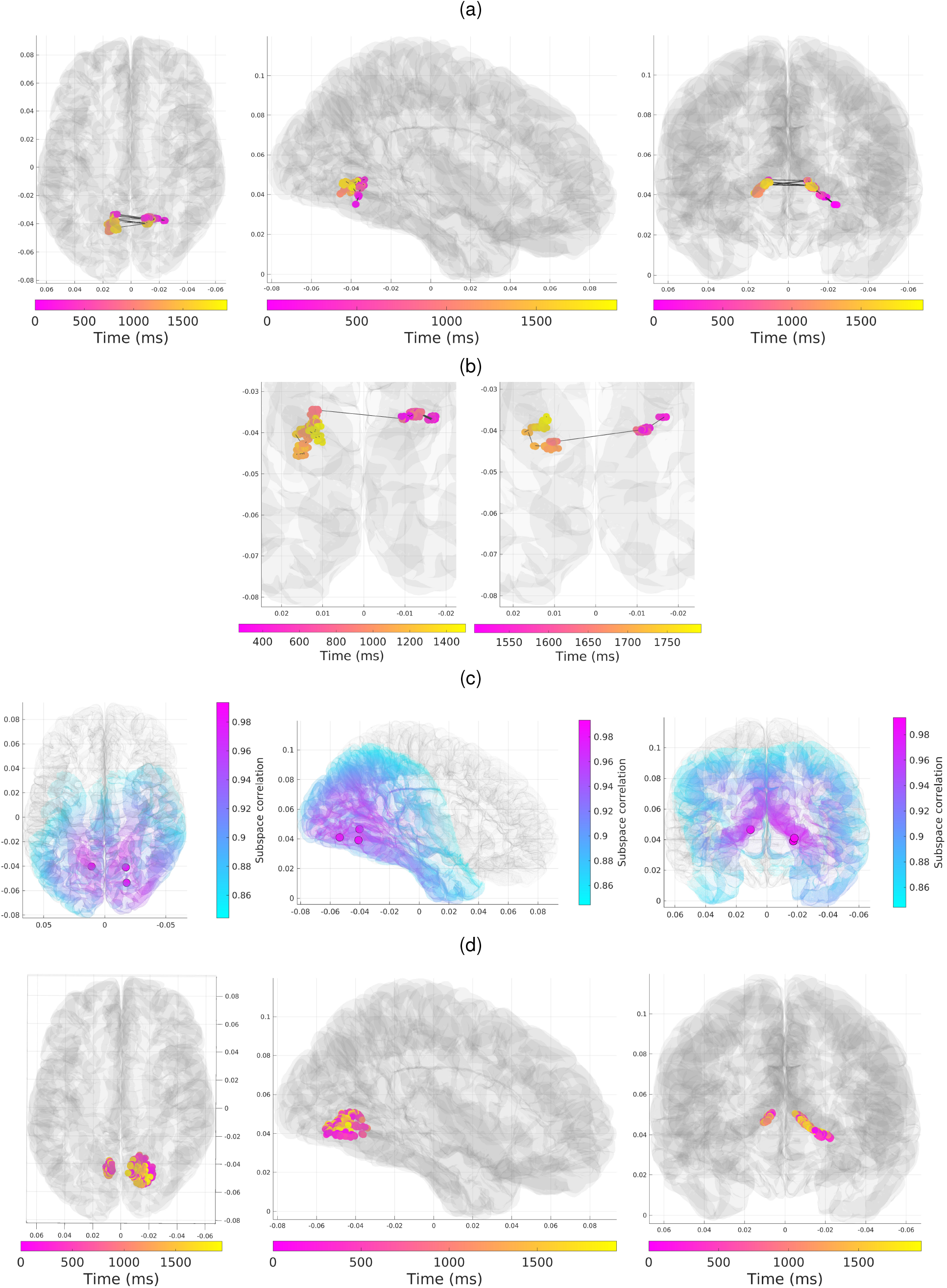
Source localization for real EEG data. **a, b)** Results of the proposed UKF-Inv method. The reconstructed trajectory of the alpha-band source is presented in full-brain views **(a)** and a zoomed-in superior view **(b)**.Successive locations are connected to visually illustrate the source propagation. **c)** Localization results using the RAP-MUSIC algorithm, which identifies three distinct static dipoles. The reported subspace correlation value corresponds to the first extracted dipole only. **d)** Results of the traditional dipole-fitting approach, where a single equivalent current dipole is estimated at each independent time slice. Across panels **a, c**, and **d**, the cortical mesh is displayed from superior, lateral, and posterior perspectives to provide a comprehensive view. For the spatiotemporal tracking methods (**a, b**, and **d**), each point represents the estimated source location at a specific time instant, with the color encoding the temporal progression in milliseconds.

Panel 15 a shows the estimated electrical activity, which exhibits a center frequency of 10.25 Hz, falling within the alpha band. Panel (b) presents the corresponding spatial coefficients utilized to track the evolving topography. The resulting source trajectory, computed by applying the subspace correlation criterion (Eq. (13)) to the estimated dynamical topography, is illustrated in Figures 16a, b. In contrast to the predominantly unilateral MEG results, the EEG activity exhibits pronounced bilateral patterns across the left and right occipital regions, which can be anatomically associated with the calcarine sulcus — a well-known generator of the alpha rhythm. The reconstructed trajectory demonstrates a physiologically plausible location alongside a highly spatiotemporally cohesive structure. A detailed close-up of this dynamic is provided in the zoomed-in superior perspectives of panel (b). These sequential temporal windows capture the source smoothly traveling across one hemisphere before rapidly transitioning to the contralateral hemisphere, where it resumes a similarly smooth propagation pattern. We speculate that this alternating interhemispheric jump may be a consequence of a transient refractory period within the initially engaged neural population, prompting the subsequent recruitment of the homologous contralateral region.

To validate these spatial findings, we applied the RAP-MUSIC method to the same EEG segment after band-pass filtering in the 8–12 Hz range, utilizing all available channels. The resulting dipole configuration is shown in Figure 16c. Consistent with the MEG analysis, RAP-MUSIC (using a subspace correlation threshold of 0.85) recovered three static dipoles located compatible with alpha-related cortical sources and lying close to the trajectory obtained via our dynamic spatiotemporal tracking model.

We further performed traditional dipole fitting independently at each time point using the same full-channel data. Although this framewise approach identifies spatially compact clusters within the occipital regions (Figure 16d), the temporal consistency is compromised. This erratic jumping behavior underscores the inherent difficulty of obtaining a coherent spatiotemporal trajectory using frame-by-frame dipole fitting alone, further highlighting the regularizing advantage of the proposed dynamic tracking procedure.

Finally, we assessed the reconstruction of the sensor-level signal. By substituting the estimated state variables back into the observation model (4) (excluding the noise term), we reconstructed the sensor time series. As shown for a single representative sensor in Figure 15c, the model explains 74% of the variance (*R*^2^ = 0.74) in the data. Analogous to the MEG findings, an ablation test substituting the dynamic spatial coefficients with their temporal averages reduces the explained variance to 64% (*R*^2^ = 0.64), reiterating the significant role of tracking continuous spatial variations. Regarding the remaining unexplained variance, higher-frequency components are inherently suppressed due to the smoothing behavior of the Kalman filtering process, which is optimized for tracking dominant oscillatory bands rather than broadband noise.

## 4 Discussion

In this work, we addressed the long-standing challenge of reconstructing spatiotemporally non-separable cortical activity from non-invasive MEG and EEG recordings. The traditional inverse problem solutions, such as minimum norm estimation [21] and dipole fitting [19, 54], operate under the assumption of spatiotemporal separability [2], which compromises their ability to track the continuous spatial migration defining cortical traveling waves [5]. To cope with this limitation, we introduced a dynamic state-space framework that explicitly decouples the rhythmic electrical dynamics of a cortical source from the slow migration of its spatial topography. By casting the source topography as a time-varying combination of data-driven eigen-topographies, we reformulated the inverse problem within the Unscented Kalman Filtering (UKF) framework [31, 32]. This allowed us to recursively estimate both the narrowband electrical timeseries and the trajectory of the underlying source, yielding trajectory estimates that are temporally coherent by construction rather than being reconstructed independently at each sample.

Conventional techniques reconstruct each time sample independently, systematically ignoring the temporal continuity of the source trajectory. In our simulation studies, where we explicitly incorporated the temporally evolving spatial components, we proved that the proposed UKF-based inversion achieved a median correlation of 0.942 with the ground-truth electrical component, outperforming both conventional MNE (0.80) and the linear Kalman filter (0.89), which essentially represents an ablated version of our algorithm. Additionally, this framework successfully bridged the gap to empirical data, recovering the principal temporal structure of real alpha rhythms and explaining 80% (*R*^2^ = 0.80) and 74% (*R*^2^ = 0.74) of the sensor-level variance in MEG and EEG recordings, respectively. Crucially, in the ablation test where the dynamic spatial coefficients were replaced by their temporal averages, the explained variance dropped in both modalities (to 73% and 64%, respectively). These metrics support our central hypothesis under the conditions tested: the evolving source topography measurably shapes the sensor-level signals, and explicitly modeling its non-stationary structure improves the estimate of the electrical component itself. We note that the linear Kalman filter is the appropriate ablation for this particular claim, since it isolates the contribution of the dynamic topography while holding the temporal prior fixed. The comparison against frame-by-frame dipole fitting reported in Section 3.1.2 is of a different character, as it does not carry a temporal smoothness prior.

Our use of data-driven eigen-topographies to constrain the spatial dimensionality of the inverse problem shares its conceptual roots with methods like Harmony [55], which utilize spherical harmonics as a global spatial basis. Recently, this anatomy-driven approach has evolved to utilize geometric basis functions (GBFs) — Laplace-Beltrami eigenmodes computed directly on the folded cortical manifold — to track large-scale traveling waves and represent whole-brain dynamics [56]. While geometric eigenmodes elegantly capture structurally constrained wave propagation, they rely heavily on accurate individual cortical meshes and uniform wave behavior. In contrast, our UKF framework extracts the spatial subspace directly from the sensor-level data via SVD. This data-driven strategy bypasses the need for idealized anatomical harmonics, allowing the model to track idiosyncratic or highly localized propagating trajectories that might otherwise require an impractically large number of fixed geometric basis functions to represent.

When tracking propagating activity, a critical methodological concern is the inherent vulnerability of spatial smoothness priors. An overly restrictive prior may mathematically fabricate the appearance of a smooth traveling wave even when the underlying neural generators are stationary or unorganized. Our formulation counteracts this by utilizing a weak first-order autoregressive prior as a form of dynamic regularization while ensuring the UKF update step remains strongly data-driven. The robustness of this balance was examined in our two-dipole scenario. When presented with two stationary but phase-locked coherent sources — a confound known to generate illusory wave patterns in traditional static models [26] — our model successfully avoided inferring spurious propagation. Instead of artificially fabricating a continuous wave-like trajectory, the model locked onto the dominant source and maintained globally flat spatial dynamics, yielding a compact spatially stationary cluster. Recent forward-modeling approaches have shown that sensor-level phase relationships contain sufficient information to separate traveling waves from standing waves when the spatiotemporal geometry is modeled explicitly [57]. Our observation is consistent with this. We stress, however, that it rests on a single source configuration, at one dipole separation, one phase lag, and one noise level, and that the wave-versus-static judgment was made qualitatively by inspecting whether the estimated spatial coefficients drift. Turning this into a usable diagnostic would require a scalar statistic summarizing the displacement of the estimated topography, a null distribution for it, and an operating characteristic measured across dipole separations, phase lags, and signal-to-noise ratios.

The physiological plausibility of the recovered spatiotemporal trajectories further corroborates the validity of our method. When applied to real resting-state recordings, our algorithm reconstructed an alpha-band source originating near the calcarine sulcus — a canonical generator of the alpha rhythm. Interestingly, the recovered trajectory exhibited smooth propagation across one hemisphere followed by a rapid transition to the contralateral homologous region. This dynamic behavior aligns tightly with current neurobiological literature, which documents that alpha oscillations are not purely standing waves but exhibit organized intra-cortical propagation [16] and global traveling wave properties [14]. One possible interpretation of the observed interhemispheric transition is a transient refractory period within the initially engaged population, prompting subsequent recruitment of the contralateral region — a mechanism that has been linked to cross-hemispheric coordination and the organization of higher-frequency bursts [58, 13]. We deliberately refrain from committing to this interpretation. A competing and equally consistent explanation is that two bilaterally distributed, coherent alpha generators are present and that our single-source model, fitted to their mixture, produces an apparent source that alternates between hemispheres. However, even within each hemisphere, we did observe spatial displacements captured by our model.

### 4.1 Limitations and scope

The following limitations and practical considerations must be acknowledged.

#### Identifiability

The core observation model is bilinear (the product of an electrical scalar and spatial coefficients), which introduces an intrinsic scale ambiguity. However, rather than artificially normalizing the variables to enforce identifiability, we avoid normalization and rely on the smoothness constraints implicitly imposed by the autoregressive model on the spatial coefficients. Moreover, because the final spatial localization step evaluates the shape of the reconstructed topography, this scalar ambiguity does not affect the identified source locations. Second, our model assumes a separation of timescales, where the spatial topography evolves more slowly than the carrier oscillation. The autoregressive parameters governing this separation are not fixed a priori: they are estimated from each data segment by maximizing the marginal likelihood, as described in Section 2.6. We note, however, that the scaling coefficient *c*_**A**_ and the process noise covariance **B** both act on the smoothness of the spatial trajectory and may therefore be only weakly separately identifiable. We have not performed a profile-likelihood analysis to check this, and individual parameter values should accordingly be interpreted with caution even though the resulting state estimates are stable across our 1000 simulation runs.

#### Propagation velocity

The method requires the topography to migrate more slowly than the carrier oscillation, and Table 2 delimits the resulting operating envelope. Estimation is stable for envelope speeds up to approximately 0.1 m/s and degrades progressively beyond it: at 0.5 m/s the median correlation of the leading spatial coefficient falls to 0.05 and the time-averaged localization error grows to 4.4 mm. The approach is therefore suited to the slow, locally propagating regime, and trajectories recovered from faster mesoscopic waves should be interpreted with corresponding caution. Two aspects of this characterization remain open: the parameter sweep was performed at a single noise level and on paths within the calcarine sulcus, leaving it unknown how the envelope shifts with varying signal-to-noise ratios under different velocities, and how it is influenced by cortical geometry.

#### Noise characteristics

While we characterized the algorithm’s robustness across a range of signal-to-noise ratios, the simulations employed purely additive uncorrelated white noise. The impact of realistic neurophysiological background activity, which is typically spatially and temporally correlated, remains to be systematically investigated.

#### Sample size in the empirical analyses

The MEG and EEG results each derive from a single participant and a single selected segment containing a pronounced alpha spindle. These serve to demonstrate that the method runs on real recordings and returns anatomically sensible output. They do not support inferences about alpha propagation as a population-level phenomenon, and we have not assessed reproducibility across segments within a participant or across participants.

#### Anatomical realizability of the estimated topography

The latent topography 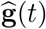 evolves freely within the SVD subspace and is not constrained to lie on the manifold of physically realizable leadfield topographies; anatomy enters only at the subsequent localization step. Reporting the attained subspace correlation as a function of time would indicate how far the estimated topography strays from that manifold, and a formulation that parameterizes the source position directly and propagates it on the cortical mesh would remove the issue altogether at a considerably greater computational cost.

#### Choice of estimator

The observation model is bilinear and, hence, conditionally linear in each block of the state. Conditionally linear or Rao–Blackwellized filtering schemes are therefore applicable and might be both more accurate and cheaper than the unscented transform. We adopted the UKF for its generality and did not benchmark it against such alternatives.

#### Baseline comparisons

As noted above, the dipole-fitting comparison is not smoothness-matched. In addition, the minimum-norm estimate was summarized by averaging across the vertices of the region of interest, which attenuates the estimate when dipole orientations vary within the region; an orientation-aware summary would give minimum-norm estimation a fairer hearing.

Finally, our current analysis focused on scenarios dominated by a single primary traveling wave. While the model separated a genuine traveling wave from a spurious wave-like sensor pattern in the one configuration we tested, resolving multiple simultaneously propagating and overlapping waves represents a substantially more complex challenge that we have not attempted. Nevertheless, our framework seamlessly integrates with existing discrete-source algorithms. We advocate a pipeline approach: leveraging our dynamic spatial tracking to diagnose whether an observed pattern stems from a propagating wave, and deploying methods from the MUSIC family [54, 34] to extract specific static components. By accommodating the spatiotemporal inseparability of brain activity within the inverse model itself, we hope this approach offers a useful starting point for the non-invasive investigation of macro-scale brain dynamics in cognitive neuroscience — where traveling waves are increasingly implicated in sensory gating and perception [9, 10] — and, subject to the validation outlined above, in clinical settings such as the tracking of interictal epileptiform discharges [43].

## 5 Conclusion

Non-invasive mapping of cortical traveling waves is fundamentally challenged by the spatiotemporal mixing of underlying neural dynamics. To address this, we introduced a novel analytical approach based on the Unscented Kalman Filter that explicitly models the EEG/MEG signal as a bilinear combination of a narrowband stochastic electrical oscillator and a slowly varying spatial topography within a data-driven SVD subspace. Evaluated on a large-scale simulation scenario, the approach recovered the electrical time course more accurately than minimum norm estimation and an ablated linear Kalman filter, and produced more temporally coherent trajectories than frame-by-frame dipole fitting. Additionally, when testing our method across varying SNR levels and simulated envelope propagation velocities from 0.01 to 0.5 m/s, we observed that trajectory reconstruction is accurate up to approximately 0.1 m/s and degrades at higher speeds, as the timescale separation underlying the model is progressively violated. In a controlled two-dipole simulation, it did not fabricate propagation where none existed. Reproducing these properties across participants remains to be done; within the slow propagation regime characterized here, we believe the approach offers a practical route to the non-invasive study of macroscale propagating cortical dynamics.

## Ethics statement

The studies involving humans were approved by the HSE University Committee on Inter-University Surveys and Ethical Assessment of Empirical Research. The studies were conducted in accordance with local legislation and institutional requirements. The participant provided your written informed consent to participate in this study.

The study used publicly available MEG data [35] whose collection was approved by the NIH Institutional Review Board (Recruitment and Characterization of Healthy Research Volunteer for NIMH Intramural Studies NCT033046). The authors of the current manuscript declare no competing interests.

## Acknowledgments

This work is an output of a research project HSE-BR-2025-26 implemented as part of the Basic Research Program at HSE University.

## Appendix

## Subspace Correlation

Our goal is to compute the subspace correlation metric between the model subspace spanned by a local leadfield matrix **G** (*r*) and the estimated topography 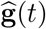, which is reconstructed from the spatial coefficients for each time point *t* and each cortical vertex *r*:

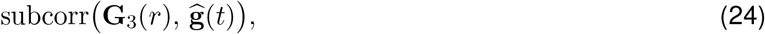

where 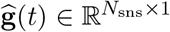 and 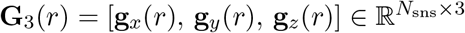.

Recall that **g**_*x*_(*r*), **g**_*y*_(*r*), and **g**_*z*_(*r*) *r* denote the topographies of a unit dipole placed at vertex and oriented along the *x*-, *y*-, and *z*-axes, respectively. Thus, the objective is to evaluate the principal angle (subspace correlation) between the column subspace of **G** (*r*) and the vector 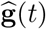.

Below, we outline the matrix formulation used to compute the subspace correlation across a large number of vertices (ranging from ≈1300 to ≈270,000 for the different cortical meshes used in this study).

First, for each vertex *r*, we compute the upper triangular matrix **R**(*r*) via the Cholesky decomposition of the matrix **A**(*r*) = **G**_3_(*r*)^*T*^ **G**_3_(*r*), where **A**(*r*) ∈ ℝ^3*×*3^. This decomposition needs to be computed only once and stored for subsequent steps.

Next, we construct the matrix 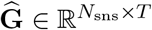, whose columns are the normalized topography vectors 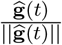.

Then, for each vertex, we compute the following matrix **Z**:

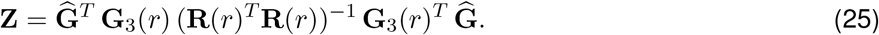

The square roots of the diagonal elements of this matrix **Z** yield the desired subspace correlations for each time point *t*.

Finally, to compute the subspace correlation for the conventional dipole fitting task, we simply replace the estimated topography 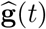 with the corresponding sensor data vector **y**(*t*).

## Modeling Leadfield Errors

Leadfield errors are modeled by perturbing each column of the gain matrix according to a Gaussian probability distribution.

The radial perturbations are represented as independent univariate Gaussian distributions aligned with the directions of the leadfield vectors. These errors are assumed to be proportional to the Euclidean norm of each leadfield column. Specifically, the *k*-th column of **G**_rad_ is defined as *β*_rad_ *ξ* ||**g**_*k*_|| **g**_*k*_, where **g**_*k*_ denotes the *k*-th column of **G** and ||**g**_*k*_|| is its Euclidean norm. The term *ξ* follows a standard univariate Gaussian distribution, while the parameter *β*_rad_ determines the strength of the radial errors. This type of perturbation essentially modifies the magnitude of the original leadfield vector, thereby introducing uncertainty into the gain amplitude. Such inaccuracies may arise from errors in estimating tissue conductivity or sensor distance, rather than discrepancies in orientation geometry.

In contrast, tangential perturbations are designed to alter the direction of the leadfield vector while preserving its original magnitude. This effectively simulates errors in dipole orientation or sensor alignment. The tangential perturbation to the *k*-th leadfield is obtained by 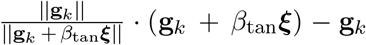, where ***ξ*** follows an *N*_sns_ -dimensional standard Gaussian distribution, and *β*_tan_ scales the intensity of the orientation error.

The parameters *β*_rad_ and *β*_tan_ were selected to ensure that the total additive perturbation reached approximately 10% of the standard deviation of the original gain matrix elements. Furthermore, the tangential error was intentionally constrained to be significantly smaller than the radial component. Given that the leadfield coefficients are on the order of 10^−5^, we set *β*_rad_ = 2.5 *×* 10^−4^ and *β*_tan_ = 2.5 *×* 10^−6^.

## Footnotes

1 The coefficient of determination, denoted as *R*^2^, is defined as 1 minus the ratio of the residual variance to the total variance. It indicates the proportion of the original signal’s variance that is captured by the model.

## References

[1] Franciscus Cornelis Donders. On the speed of mental processes. Acta psychologica, 30:412–431, 1969.

[2] David M Alexander, Chris Trengove, and Cees van Leeuwen. Donders is dead: cortical traveling waves and the limits of mental chronometry in cognitive neuroscience. Cognitive Processing, 16:365–375, 2015.

[3] Martin Seeber, Lucia-Manuela Cantonas, Mauritius Hoevels, Thibaut Sesia, Veerle Visser-Vandewalle, and Christoph M Michel. Subcortical electrophysiological activity is detectable with high-density eeg source imaging. Nature communications, 10(1):753, 2019.

[4] Sylvain Baillet. Magnetoencephalography for brain electrophysiology and imaging. Nature neuroscience, 20(3):327–339, 2017.

[5] Lyle Muller, Frédéric Chavane, John Reynolds, and Terrence J Sejnowski. Cortical travelling waves: mechanisms and computational principles. Nature Reviews Neuroscience, 19(5):255–268, 2018.

[6] David M Alexander, Andrey R Nikolaev, Peter Jurica, Mikhail Zvyagintsev, Klaus Mathiak, and Cees van Leeuwen. Global neuromagnetic cortical fields have non-zero velocity. PLoS One, 11(3):e0148413, 2016.

[7] Justin M Campbell, Tyler S Davis, Daria Nesterovich Anderson, Amir Arain, Zachary W Davis, Cory S Inman, Elliot H Smith, and John D Rolston. Macroscale traveling waves evoked by single-pulse stimulation of the human brain. Journal of Neuroscience, 45(21), 2025.

[8] David M Alexander, Tonio Ball, Andreas Schulze-Bonhage, and Cees van Leeuwen. Large-scale cortical travelling waves predict localized future cortical signals. PLoS computational biology, 15(11):e1007316, 2019.

[9] Zachary W Davis, Lyle Muller, Julio Martinez-Trujillo, Terrence Sejnowski, and John H Reynolds. Spontaneous travelling cortical waves gate perception in behaving primates. Nature, 587(7834):432–436, 2020.

[10] Sayak Bhattacharya, Scott L Brincat, Mikael Lundqvist, and Earl K Miller. Traveling waves in the prefrontal cortex during working memory. PLoS computational biology, 18(1):e1009827, 2022.

[11] Lyle Muller, Giovanni Piantoni, Dominik Koller, Sydney S Cash, Eric Halgren, and Terrence J Sejnowski. Rotating waves during human sleep spindles organize global patterns of activity that repeat precisely through the night. Elife, 5:e17267, 2016.

[12] Zachary J Haigh, Harry Tran, Taylor Berger, Sina Shirinpour, Ivan Alekseichuk, Seth Koenig, Jan Zimmermann, Robert McGovern, David Darrow, Alexander Herman, et al. Modulation of motor excitability reflects traveling waves of neural oscillations. Cell reports, 44(6), 2025.

[13] Uma R Mohan, Honghui Zhang, Bard Ermentrout, and Joshua Jacobs. The direction of theta and alpha travelling waves modulates human memory processing. Nature Human Behaviour, 8(6):1124–1135, 2024.

[14] Honghui Zhang, Andrew J Watrous, Ansh Patel, and Joshua Jacobs. Theta and alpha oscillations are traveling waves in the human neocortex. Neuron, 98(6):1269–1281, 2018.

[15] Adeeti Aggarwal, Connor Brennan, Jennifer Luo, Helen Chung, Diego Contreras, Max B Kelz, and Alex Proekt. Visual evoked feedforward-feedback traveling waves organize neural activity across the cortical hierarchy in mice. Nature communications, 13(1):4754, 2022.

[16] Rikkert Hindriks, Michel JAM van Putten, and Gustavo Deco. Intra-cortical propagation of eeg alpha oscillations. Neuroimage, 103:444–453, 2014.

[17] Sarah F Schoch, Brady A Riedner, Sean C Deoni, Reto Huber, Monique K LeBourgeois, and Salome Kurth. Across-night dynamics in traveling sleep slow waves throughout childhood. Sleep, 41(11):zsy165, 2018.

[18] Andrey Andreevich Gorshkov, Alexei Evgenievich Ossadtchi, and Alexander Lvovich Fradkov. Regularization of the eeg/meg inverse problem by a local cortical wave pattern. Information and Control Systems, pages 12–20, 2017.

[19] Michael Scherg et al. Fundamentals of dipole source potential analysis. Auditory evoked magnetic fields and electric potentials. Advances in audiology, 6(40–69):25, 1990.

[20] John C Mosher and Richard M Leahy. Recursive music: a framework for eeg and meg source localization. IEEE transactions on biomedical engineering, 45(11):1342–1354, 1998.

[21] Matti S Hamalainen. Interpreting measured magnetic fields of the brain: estimates of current distributions. Helsinki Univ. of Technol., Rep, 1984.

[22] Britta U Westner, Sarang S Dalal, Alexandre Gramfort, Vladimir Litvak, John C Mosher, Robert Oostenveld, and Jan-Mathijs Schoffelen. A unified view on beamformers for m/eeg source reconstruction. NeuroImage, 246:118789, 2022.

[23] Matthew J Brookes, James Leggett, Molly Rea, Ryan M Hill, Niall Holmes, Elena Boto, and Richard Bowtell. Magnetoencephalography with optically pumped magnetometers (opm-meg): the next generation of functional neuroimaging. Trends in Neurosciences, 45(8):621–634, 2022.

[24] Joonas Iivanainen, Matti Stenroos, and Lauri Parkkonen. Measuring meg closer to the brain: Performance of on-scalp sensor arrays. NeuroImage, 147:542–553, 2017.

[25] Leonhard Schreiner, Michael Jordan, Sebastian Sieghartsleitner, Christoph Kapeller, Harald Pretl, Kyousuke Kamada, Priscella Asman, Nuri F Ince, Kai J Miller, and Christoph Guger. Mapping of the central sulcus using non-invasive ultra-high-density brain recordings. Scientific reports, 14(1):6527, 2024.

[26] Alexander Zhigalov and Ole Jensen. Perceptual echoes as travelling waves may arise from two discrete neuronal sources. NeuroImage, 272:120047, 2023.

[27] Camilo Lamus, Matti S Hä mäläinen Simona Temereanca, Emery N Brown, and Patrick L Purdon. A spatiotemporal dynamic distributed solution to the meg inverse problem. NeuroImage, 63(2):894–909, 2012.

[28] Joonas Lahtinen, Paavo Ronni, Narayan Puthanmadam Subramaniyam, Alexandra Koulouri, Carsten Wolters, and Sampsa Pursiainen. Standardized kalman filtering for dynamical source localization of concurrent subcortical and cortical brain activity. Clinical Neurophysiology, 168:15–24, 2024.

[29] Takeru Matsuda and Fumiyasu Komaki. Time series decomposition into oscillation components and phase estimation. Neural Computation, 29(2):332–367, 2017.

[30] Simon J Julier and Jeffrey K Uhlmann. New extension of the kalman filter to nonlinear systems. In Signal processing, sensor fusion, and target recognition VI, volume 3068, pages 182–193. Spie, 1997.

[31] Simon J Julier and Jeffrey K Uhlmann. Unscented filtering and nonlinear estimation. Proceedings of the IEEE, 92(3):401–422, 2004.

[32] Eric A Wan and Rudolph Van Der Merwe. The unscented kalman filter. Kalman filtering and neural networks, pages 221–280, 2001.

[33] John C Mosher and Richard M Leahy. Eeg and meg source localization using recursively applied (rap) music. In Conference Record of The Thirtieth Asilomar Conference on Signals, Systems and Computers, pages 1201–1207. IEEE, 1996.

[34] John C Mosher and Richard M Leahy. Source localization using recursively applied and projected (rap) music. IEEE Transactions on signal processing, 47(2):332–340, 1999.

[35] Allison C Nugent, Adam G Thomas, Margaret Mahoney, Alison Gibbons, Jarrod T Smith, Antoinette J Charles, Jacob S Shaw, Jeffrey D Stout, Anna M Namyst, Arshitha Basavaraj, et al. The nimh intramural healthy volunteer dataset: A comprehensive meg, mri, and behavioral resource. Scientific Data, 9(1):518, 2022.

[36] Risto J Ilmoniemi and Jukka Sarvas. Brain signals: physics and mathematics of MEG and EEG. Mit Press, 2019.

[37] Norbert Wiener. Nonlinear problems in random theory. MIT Press, 1966.

[38] Rudolph Emil Kalman. A new approach to linear filtering and prediction problems. Journal of Basic Engineering, 1960.

[39] Arnaud Doucet, Nando De Freitas, Neil James Gordon, et al. Sequential Monte Carlo methods in practice, volume 1. Springer, 2001.

[40] François Tadel, Sylvain Baillet, John C Mosher, Dimitrios Pantazis, and Richard M Leahy. Brainstorm: A user-friendly application for meg/eeg analysis. Computational intelligence and neuroscience, 2011(1): 879716, 2011.

[41] MX Huang, John C Mosher, and RM Leahy. A sensor-weighted overlapping-sphere head model and exhaustive head model comparison for meg. Physics in Medicine & Biology, 44(2):423, 1999.

[42] The MathWorks Inc. Matlab version: 9.14.0 (r2023a), 2023. URL https://www.mathworks.com.

[43] AA Kuznetsova and AE Ossadtchi. Analysis of the local dynamics of interictal discharge propagation using a traveling wave model. Neuroscience and Behavioral Physiology, 52(9):1436–1447, 2022.

[44] VM Verkhliutov. A model of the structure of the dipole source of the alpha rhythm in the human visual cortex. Zhurnal Vysshei Nervnoi Deiatelnosti Imeni IP Pavlova, 46(3):496–503, 1996.

[45] E. Larson, A. Gramfort, D. A. Engemann, J. Leppakangas, C. Brodbeck, M. Jas, T. L. Brooks, J. Sassenhagen, D. McCloy, M. Luessi, J.-R. King, R. Hö chenberger R. Goj, C. Brunner, G. Favelier, M. van Vliet, M. Wronkiewicz, A. Rockhill, C. Holdgraf, …, and luzpaz. Mne-python (v1.9.0), 2024. URL 10.5281/zenodo.592483.

[46] Alexandre Gramfort, Martin Luessi, Eric Larson, Denis A Engemann, Daniel Strohmeier, Christian Brodbeck, Roman Goj, Mainak Jas, Teon Brooks, Lauri Parkkonen, et al. Meg and eeg data analysis with mne-python. Frontiers in Neuroinformatics, 7:267, 2013.

[47] Pierre Comon. Independent component analysis, a new concept? Signal processing, 36(3):287–314, 1994.

[48] Vadim V Nikulin, Guido Nolte, and Gabriel Curio. A novel method for reliable and fast extraction of neuronal eeg/meg oscillations on the basis of spatio-spectral decomposition. NeuroImage, 55(4):1528–1535, 2011.

[49] Bruce Fischl. Freesurfer. Neuroimage, 62(2):774–781, 2012.

[50] Jan Kybic, Maureen Clerc, Touffic Abboud, Olivier Faugeras, Renaud Keriven, and Théodore Papadopoulo. A common formalism for the integral formulations of the forward eeg problem. IEEE transactions on medical imaging, 24(1):12–28, 2005.

[51] Alexandre Gramfort, Théodore Papadopoulo, Emmanuel Olivi, and Maureen Clerc. Openmeeg: opensource software for quasistatic bioelectromagnetics. Biomedical engineering online, 9:1–20, 2010.

[52] Arkadi Nemirovski. Interior point polynomial time methods in convex programming. Lecture notes, 42(16): 3215–3224, 2004.

[53] Per Christian Hansen. The l-curve and its use in the numerical treatment of inverse problems. InviteComputational Inverse Problems in Electrocardiology, 2000.

[54] John C Mosher, Paul S Lewis, and Richard M Leahy. Multiple dipole modeling and localization from spatiotemporal meg data. IEEE transactions on biomedical engineering, 39(6):541–557, 1992.

[55] Yury Petrov. Harmony: Eeg/meg linear inverse source reconstruction in the anatomical basis of spherical harmonics. PloS one, 7(10):e44439, 2012.

[56] James C Pang, Kevin M Aquino, Marianne Oldehinkel, Peter A Robinson, Ben D Fulcher, Michael Breakspear, and Alex Fornito. Geometric constraints on human brain function. Nature, 618(7965):566–574, 2023.

[57] Laetitia Grabot, Gwendal Merholz, Jonathan Winawer, David J Heeger, and Laura Dugué. Traveling waves in the human visual cortex: An meg-eeg model-based approach. PLOS Computational Biology, 21(4): e1013007, 2025.

[58] Ali Bahramisharif, Marcel AJ van Gerven, Erik J Aarnoutse, Manuel R Mercier, Theodore H Schwartz, John J Foxe, Nick F Ramsey, and Ole Jensen. Propagating neocortical gamma bursts are coordinated by traveling alpha waves. Journal of Neuroscience, 33(48):18849–18854, 2013.

